# Influence of age on functional memory T cell diversity

**DOI:** 10.1101/2021.05.29.446296

**Authors:** Fengqin Fang, Wenqiang Cao, Weikang Zhu, Nora Lam, Lingjie Li, Sadhana Gaddam, Yong Wang, Chulwoo Kim, Simon Lambert, Huimin Zhang, Bin Hu, Donna L.Farber, Cornelia M. Weyand, Jörg J. Goronzy

## Abstract

Memory T cells exhibit considerable diversity that determines their ability to be protective and their durability. Here, we examined whether changes in T cell heterogeneity contribute to the age-associated failure of immune memory. By screening for age-dependent T cell surface markers, we have identified CD4 and CD8 memory T cell subsets that are unrelated to previously defined subsets of central and effector memory cells. Memory T cells expressing the ecto-5’-nucleotidase CD73 constitute a functionally distinct subset of memory T cells that declines with age. They exhibit many features favorable for immune protection, including longevity and polyfunctionality. They have a low turnover, but are poised to display effector functions and to develop into cells resembling tissue-resident memory T cells (T_RM_). Upstream regulators of differential chromatin accessibility and transcriptomes include transcription factors that are characteristic for conferring these superior memory features as well as facilitating CD73 expression. CD73 is not just a surrogate marker of these regulatory networks but is directly involved in T cell survival and T_RM_ differentiation Interventions preventing the decline of this T cell subset or increasing CD73 expression have the potential to improve immune memory in older adults.

## Introduction

Immune aging is associated with a decline in immune memory. A classic example is shingles, a reactivation of the varicella zoster virus (VZV) that affects up to 50% of the human population by the age of 80 years. Examples of recurring infections are pneumococcal disease and infections with the respiratory syncytial virus, both childhood infections with increased morbidity and mortality in the elderly. Moreover, T cell immunity does not provide protection against the annual influenza infections that occur due to antigenic drifts in antibody epitopes, although T cell epitopes appear to be relatively conserved^1^. Conversely, memory cell function to other pathogens can be lastingly effective. For example, symptomatic reactivation of EBV and CMV is rarely seen with normal aging, although it occurs with immunosuppressive treatments. This dichotomy is also illustrated by the response to the live VZV vaccine Zostavax and the VZV component vaccine Shingrix that differ in their abilities to generate lasting immune memory in the elderly^2^.

Immunological memory refers to T cells that have recognized a pathogen before and generate an enhanced immune response upon re-encounter based on their increased frequencies as well as their acquired properties. Important functional domains for memory T cells are their durability, their ability to proliferate, their migration patterns and their poised state to exert effector function. Cell surface markers for conventional T cell subsets of central and effector memory (T_CM_, T_EM_) and of T_EMRA_ cells have been widely used to examine the influence of age on peripheral T cells. In general, the distribution of CD4 memory T cell subsets is relatively stable over adult life, while CD8 T_EM_ and T_EMRA_ populations accumulate. Studies of functional CD4 T cells based on their cytokine production patterns such as TH1, TH17 and TH2 T cells have not shown a consistent change with age.

Distinct from T cells primed by exogenous pathogens, naïve T cells may also acquire memory-like properties following self-antigen recognition during normal homeostasis, coined as ‘‘virtual’’ and ‘‘innate’’ memory T cells. Such virtual CD8 T cells accumulate with age. Whether they are as beneficial as normal memory cells in a recall response remains debated^3^. They lack the clonal enrichment of antigen-specific T cells, a prerequisite of immune memory, but they also have a reduced proliferative potential in old individuals^4^.

Conventional markers incompletely define the heterogeneity of memory T cells. A better understanding of functional memory cell heterogeneity has implications for vaccination strategies to improve protective and sustained immunity, including in the elderly. The recent progress in single cell cytometric and RNA-seq analysis has led to a plethora of cell surface marker combinations defining memory cell subsets that only in part have been correlated with functional properties^5,6^. To examine memory T cell heterogeneity with age, we probed a transcriptome dataset of human peripheral CD4 and CD8 T cell subsets from young and old adults for cell surface markers that changed with age (accession code: PRJNA638216). We identified CD73 as a molecule that allowed a hereto unknown subsetting of memory CD4 and CD8 T cells that does not correlate with previously defined subsets, is functionally meaningful and changes with age. CD73^+^ memory T cells excel in their durability, poised effector function and ability to differentiate into cells reminiscent of T_RM_ in vitro under TCR and subsequent TGFβ/IL-15 stimulation. Transcription factor (TF) networks that are important for memory cell function regulate transcription of CD73 that therefore identifies a selective differentiation state. Equally importantly, CD73 directly confers survival advantage in murine antiviral responses as well as the ability to persist as tissue-residing T cells expressing T_RM_-associated markers, such as CD69 and CXCR6. Unlike CD73^+^ T cells, CD73^-^ memory CD4 T cells are a heterogeneous population that increases with age and includes actively replicating, short-lived cells largely devoid of polyfunctional T cells. Identifying means to prevent the decline of CD73^+^ T cells with age has the potential to improve immune memory in the elderly.

## Results

### Age-associated changes in memory T cell heterogeneity

In RNA-sequencing studies of peripheral T cell subsets, we found that the expression of *NT5E* encoding the 5’-nucleotidase CD73 declined with age. Together with our recent finding that *ENTPD1*-encoded CD39 is more readily induced in memory T cells from older individuals^7^, these data raise the possibility that molecules involved in purinergic signaling contribute to memory cell heterogeneity.

To examine the relationship of CD73 expression to conventional T cell subset definitions, we compared peripheral blood mononuclear cells (PBMCs) from fourteen 20 to 35 and twelve 65-80 year-old healthy adults. CD73 was expressed on all CD4 and CD8 T cell subsets. For CD4 cells, expression was more frequent in naïve and T_EMRA_ cells with 25% each compared to 15% in T_CM_ (Figure 1A). Expression of CD73 in CD8 T cell subsets was generally higher, with most naïve and close to 40% of T_CM_ and T_EM_ CD8 T cells expressing CD73 (Figure 1B). CD8 T_EMRA_ cells had the lowest expression of CD73. Importantly, CD73 expression did not correlate with the expression of chemokine receptors that are generally used to subset memory T cells. For all CD4 and CD8 T cell subsets, except CD8 T_CM_, expression of CD73 significantly decreased with age (Figures 1A and 1B).

**Figure 1:**
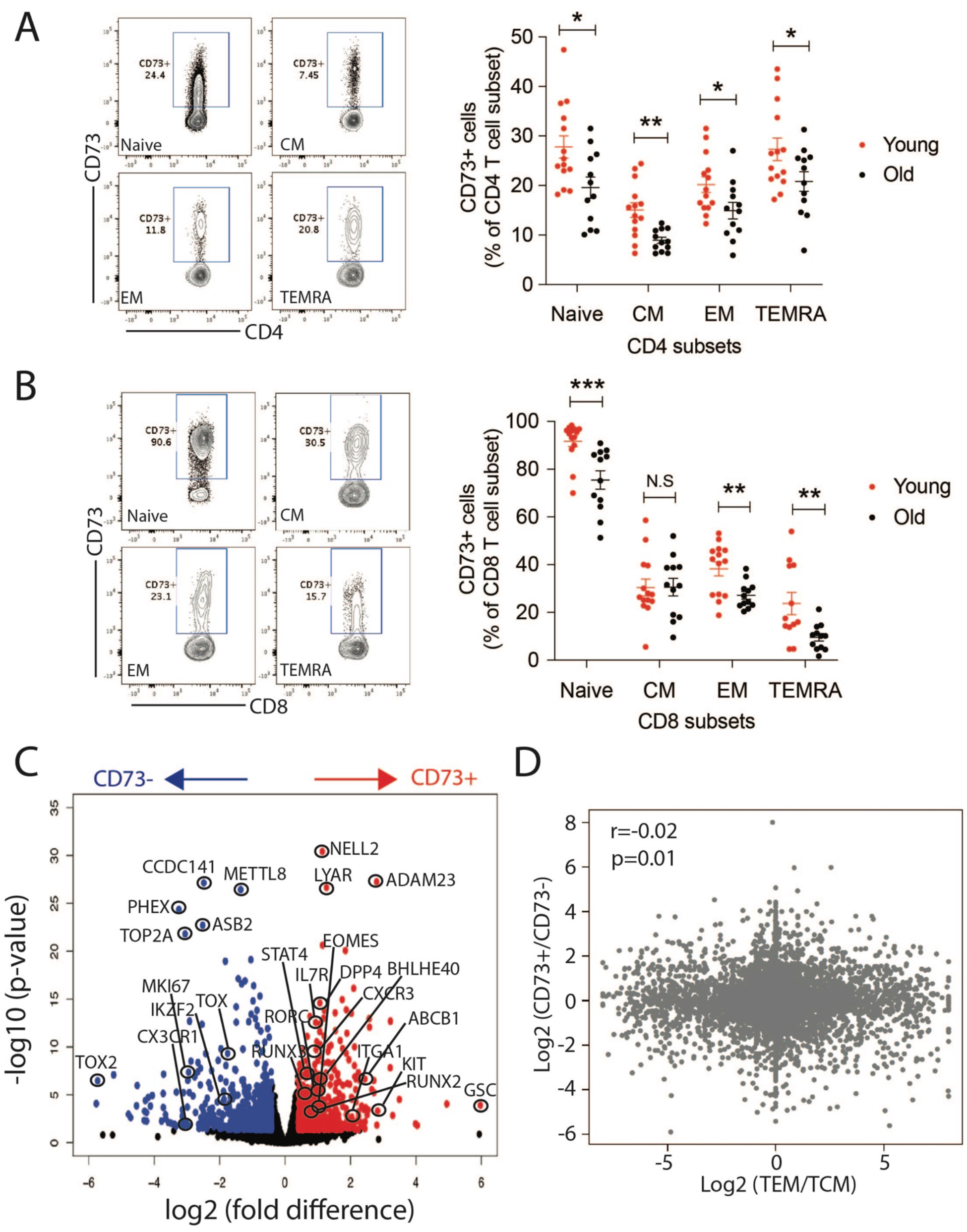
Expression of CD73 identifies a memory T cell subset that is distinct from conventional subsets and that decreases with age. **(A and B)**: CD73 expression in peripheral CD4 (A) or CD8 (B) subsets from young (<35y, red) and older (>65y, black) individuals. Results are shown as representative contour plots (left) and as dot plots of frequencies with means as indicated. Frequencies in T cells from young and old adults were compared by unpaired t-test; *p<0.05, **p<0.01, ***p<0.001. N.S: not significant. **(C)** Volcano plot of genes differentially expressed in CD73^+^ and CD73^-^ memory CD4 T cells. Genes significantly (p< 0.05) higher (red) or lower (blue) in expression by >30% in CD73^+^ T cells are indicated by color. Selected genes of immunological relevance are labeled. (D) Transcriptomes of CD73^+^ and CD73^-^ memory CD4 T cells and conventional T_CM_ and T_EM_ are compared. Results are shown as the fold difference in T_CM_ and T_EM_ plotted vs the fold difference in CD73^+^ and CD73^-^ memory CD4 T cells. RNA-seq data of T_EM_ and T_CM_ from 3 donors were downloaded from NCBI public database GSE97863 ^15^. The correlation coefficient and p value were calculated based on Pearson’s product-moment correlation analysis.

### CD73^+^ CD4^+^ T cells display a transcriptome of superior effector as well as memory cell function

To identify functional differences, we sorted human CD73^+^ and CD73^-^ memory CD4 T cells from three adults for RNA-sequencing. 634 genes were upregulated in the CD73^+^ subset, while 594 genes were downregulated as shown in the volcano plot in Figure 1C with selected genes highlighted. Genes that were more highly expressed in CD73^+^ T cells included genes characteristic of effector T cells, such as *RUNX3, RORC, PRDM1, STAT4, TBX21, IL12R, IL23R* and *CXCR3*^8^, In parallel, selected genes pertaining to long-lived memory T cells such as *IL7R, EOMES*^9^, *RUNX2*^10,11^, *ABCB1* (*MDR1*) and *CD161* (*KLRB1*)^12^ were also more highly expressed. Moreover, expression of *BHLHE40*, critical to maintaining fitness and functionality of T_RM_ and tissue-infiltrating T cells was increased^13,14^. c-KIT, also overexpressed in CD73^+^ T cells, is an additional survival factor that so far has not been implicated in memory cell longevity. To further examine the relationship of CD73^+^ cells to T_CM_ or T_EM_ cells, we retrieved RNA-seq data from the NCBI database (GSE97863) comparing human T_CM_ and T_EM_ cells^15^. We did not find a correlation of the differences between CD73^+^ and CD73^-^ cells with those between T_CM_ and T_EM_ cells, supporting the notion that subsetting memory CD4 T cells based on CD73 expression is distinct from traditional T_CM_ and T_EM_ cells.

GO term analysis, using the DAVID Bioinformatics tool, yielded a significant enrichment in CD73^+^ cells for “positive regulation of IFNγ production”, “inflammatory response”, “response to virus” and “positive regulation of cell migration” (Figure 2A). Gene set enrichment analysis (GSEA) confirmed the correlation with inflammatory response gene expression (Figure 2B). Moreover, a strong correlation of CD73-positivity was found with ribosomal gene expression (Figure 2B), reminiscent of the increased ribosomal activity in antigen-stimulated effector CD8 T cells that has been implicated in memory fate decisions^16^. Further analysis of the transcriptome data showed that CD73^+^ and CD73^-^ memory T cells, although equally represented in central and effector memory T cells, exhibited very different propensities in migratory patterns (Suppl. Figure 1A). CD73^+^ cells have decreased expression of CCR3, CCR4, CCR7, CCR8 and CXCR5, but increased transcription of CCR2, CCR6, CCR9, CXCR3, CXCR4 and CXCR6, a pattern which enables cells to migrate to peripheral tissues^17-19^ including tumors^20,21^.

**Figure 2:**
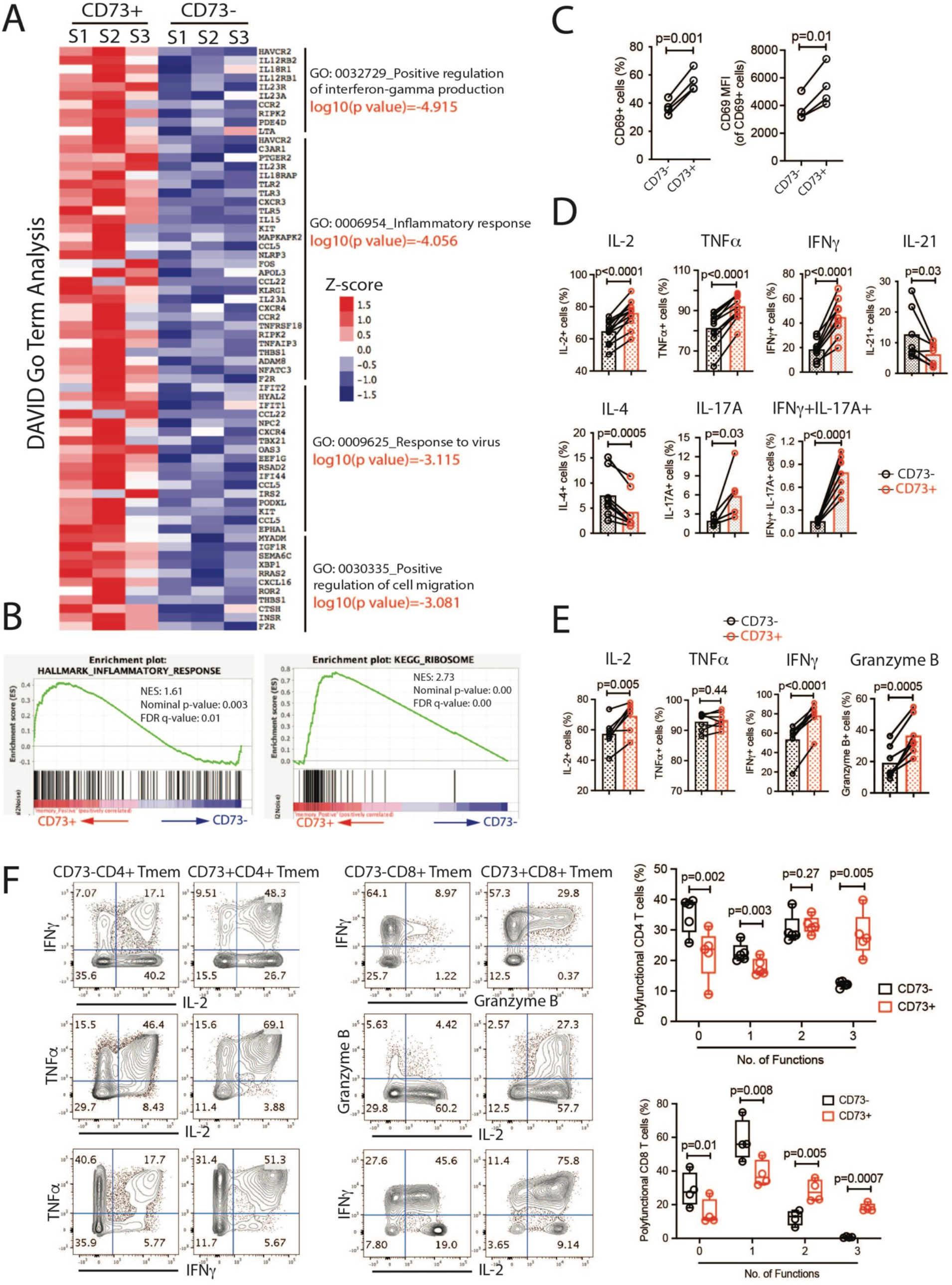
Distinctive features of the transcriptome of CD73^+^ memory CD4 T cells. **(A)** Genes overexpressed by CD73^+^ T cells were analyzed for functional enrichment using the DAVID Bioinformatics Tool. Genes from selected enriched gene ontology terms (p<0.001) are shown as heat plots. **(B)** Gene set enrichment analysis (GSEA) of transcriptome data showed a correlation for CD73^+^ cells with Hallmark_Inflammatory response and KEGG_Ribosome. (**C)** Purified CD73^+^ and CD73^-^ memory CD4 T cells were activated by plate-immobilized anti-CD3/CD28 Abs; CD69 cell surface expression was determined after 12 hours. **(D-E)** Memory CD4 (D) and CD8 (E) T cells were stimulated with PMA/ionomycin for 3-4 hours before intracellular cytokine staining. Results are shown as the frequencies of cells producing indicated cytokines. MFIs of cytokine expression is shown in Suppl. Figure 1D/E. **(F)** Data were analyzed for the co-expression of IL2, TNFα and IFNγ for CD4 T cells and IL2, granzyme B and IFNγ for CD8 T cells. Results are shown as representative contour plots and box plots of the percentage of polyfunctional T cells in the CD73^+^ (red) and CD73^-^ (black) subsets. Data were compared by two-tailed paired t-test.

Moreover, the two subsets differed in the expression of CD4 sub-lineage-defining genes^22^, CD73^+^ cells exhibited TH1, TH17 and TH1.17 gene patterns, while Treg and Th2 signatures were found in CD73^-^ T cells (Suppl. Figure 1B). However, both CD73+ and CD73^-^ T cells were diverse populations, each including different lineages. For example, Tregs defined by FOXP3 expression accounted only for a minority of less than 15% of CD73^-^ memory T cells (Suppl. Figure 1C).

To compare functional properties of CD73^+^ and CD73^-^ memory T cells, we tested their response to activation signals. CD69 was more readily induced in CD73^+^ cells upon CD3/CD28 triggering, indicating a greater responsiveness of this subset to TCR stimulation (Figure 2C). Intracellular cytokine staining upon ionomycin and PMA stimulation showed higher frequencies of CD73^+^ memory CD4 T cells that were poised to secrete IL-2, TNFα, IFNγ and/or IL-17A and fewer producers of IL-21 and IL-4 (Figure 2D). In addition, based on the MFI in the gated cytokine-positive population, the amount of IL-2, TNFα and IFNγ produced per cell was higher (Suppl. Figure 1D). Similarly, CD73^+^ memory CD8 T cells generated more TNFα per cell and were more frequently able to produce IL-2, IFNγ and granzyme B compared to CD73^-^ cells (Figure 2E and Suppl. Figure 2E). Moreover, a higher frequency of CD73^+^ memory T cells was polyfunctional. Up to 30% of CD73^+^ memory CD4 T cells secreted all three cytokines tested (IFNγ, IL-2 and TNFα) compared to 13% of CD73^-^ cells after 6 hours of anti-CD3/CD28 stimulation (Figure 2F). In contrast, the percentage of cells secreting none or only one cytokine was significantly higher in CD73^-^ memory CD4 T cells. A similar bias for polyfunctionality (here defined as the coproduction of IL-2, IFNγ and granzyme B) was observed for CD8 memory T cells (Figure 2F).

### Resistance of CD73^+^ T cells to undergo cell death

While CD73^+^ memory T cells had superior effector function, transcriptome analysis also indicated an increased expression of growth factor receptors and survival factors, unlike short-lived effector T cells (Figure 1C). Quantitative RT-PCR assays confirmed the elevated expression of *BCL2, IL7R* and *KIT* by CD73^+^ cells (Figure 3A). In vitro culture in the absence of cytokines showed a 3-fold higher propensity of CD73^-^ T cells to undergo apoptosis (Figure 3B).

**Figure 3:**
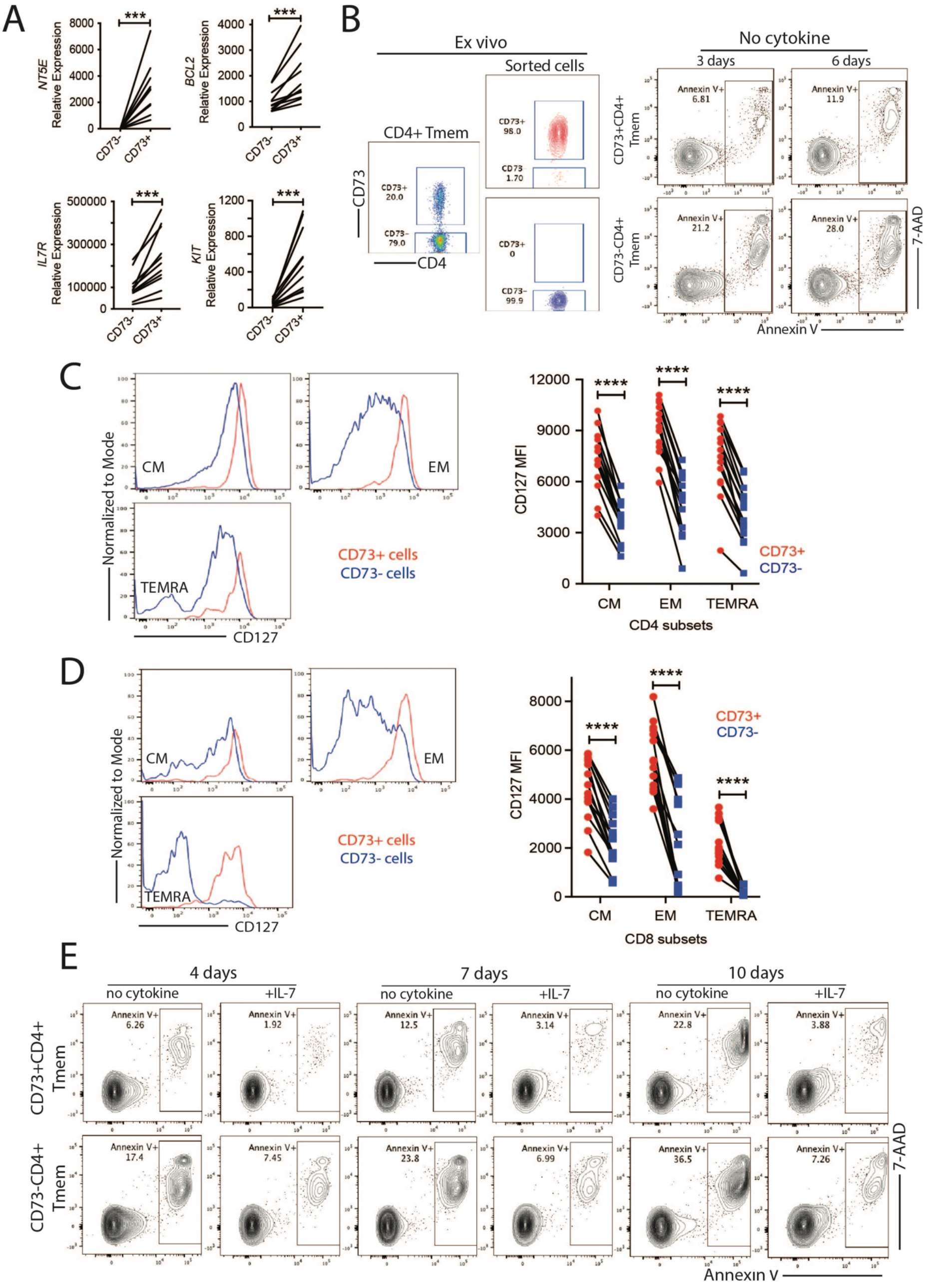
increased longevity of CD73^+^ memory T cells. **(A)** *NT5E, BCL2, IL7R* and *KIT* expression quantified by RT-PCR. Data are shown as 2^(-delta Ct)^ *10^−5^. **(B)** CD73^+^ and CD73^-^ memory CD4 T cells, respectively were cultured in the absence of cytokines. Cells were stained with Annexin V and 7-AAD. Contour plots are representative of 3 experiments. **(C and D)** CD127 (IL7R) expression on CD73^+^ vs CD73^-^ cells of CD4 (C) and CD8 (D) memory T cell subsets. Representative histograms (left) and summary data (right). **(E)** Cells were cultured in the presence or absence of IL-7 (10 ng/ml) for indicated times and stained with Annexin V and 7-AAD. Contour plots are representative of 4 experiments. Data were compared by two-tailed paired t-test. ***p<0.001, ****p<0.0001.

Differential expression of the IL7R receptor on CD73^+^ T cells was consistent for the T_CM_, T_EM_ and T_EMRA_ subsets of CD4 and CD8 memory T cells (Figure 3C and D). It is noteworthy that with exception of CD8 T_EMRA_ cells and a subset of CD8 T_EM_ cells, IL7R was still detectable on CD73^-^ cells, but clearly reduced on a per cell basis. In vitro culture in the presence of IL-7 showed that both populations dramatically increased their survival rate, however, the benefit was more pronounced for CD73^+^ T cells consistent with increased growth factor receptor expression conferring a survival advantage (Figure 3E). CD117 encoded by *c-KIT* was expressed on a small subset of CD73^+^ CD4 and CD8 memory T cells and here particularly on CD73^+^ T_EM_ cells, but was mostly absent in CD73^-^ cells of all subsets (Suppl. Fig. 2A). Given the small population size, culture with stem cell factor (SCF) treatment did not provide a global, detectable survival benefit (Suppl. Figure 2B). c-KIT expression declined with age in T_CM_ and T_EM_ subsets of CD4 and CD8 T cells (Suppl. Figure 2C).

### CD73^-^ memory CD4 T cells are a high turnover population with limited effector function

CD73^-^ T cells represent the majority of central and effector memory CD4 T cells, but their transcriptome suggested an inferior effector function. DAVID GO term analysis of the CD73^-^ cell transcriptome was enriched for the terms of “cell division” and “regulation of cell cycle” as well as “MHC class II molecules” (Figure 4A). Consistent with the DAVID analysis, GSEA showed a high correlation with the hallmark categories of “E2F targets” and “mitotic spindle” as well as with two widely-used gene sets distinguishing quiescent and dividing cells^23^ (Figure 4B).

**Figure 4:**
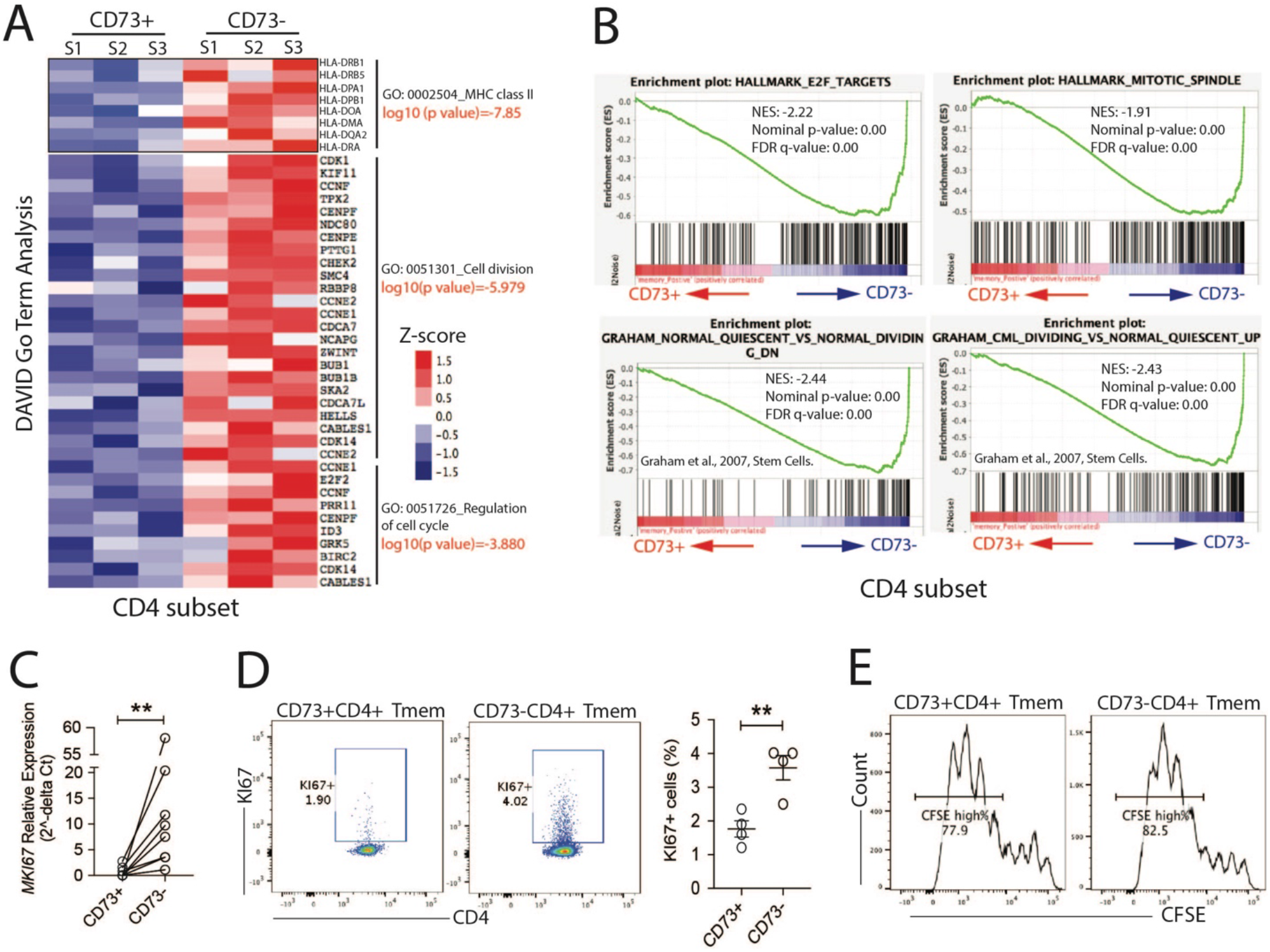
Distinctive features of the transcriptome of CD73^-^ memory CD4 T cells. **(A)** Gene ontology terms enriched in the transcriptome of CD73^-^ T memory cells indicated differences in T cell activation and cell division (p<0.001). Expression of selected genes are shown as heat plot. **(B)** GSEA shows enrichment of the CD73^-^ cell transcriptome for several hallmarks related to cell division. **(C)** *MKI67* expression in CD73^+^ and CD73^-^ memory T cells. qPCR data are shown as 2^(-delta Ct)^ *10^−5^. **(D)** Percentages of Ki67^+^ cells in CD73^+^ and CD73^-^ T cells by flow cytometry. Representative flow scatter plot (left) and mean±SEM of four experiments. **(E)** Proliferative responses of CD73^+^ and CD73^-^ memory T cells 5 days after activation with anti-CD3/CD28 Dynabeads. CFSE dilution is representative of three experiments. Results are compared by two-tailed paired t-test. **p<0.01.

To obtain further evidence for high in vivo turnover, we determined ex vivo expression of Ki67, that is restricted to actively and recently cycling cells^24^. *MKI67* mRNA was almost undetectable in CD73^+^ and clearly elevated in CD73^-^ CD4 T cells (Figure 4C). Flow cytometry detected around 4% of Ki67^+^ cells in CD73^-^ CD4 memory T cells compared to less than 2% in CD73^+^ cells (Figure 4D). The difference in cycling cells was not due to an irreversible defect; the proliferative response to anti-CD3/CD28 Ab activation was nearly equal between the two subsets (Figure 4E).

### CD73^+^ and CD73^-^ memory T cells are governed by distinct transcriptional regulatory networks

Transcriptional profiling of transcription factors that define T cell differentiation states showed that CD73^+^ T cells have increased expression of effector (including RORC, PRDM1, RUNX3 and TBX21) as well as memory cell-determining TFs (including RUNX2, EOMES and BHLHE40) compared to CD73^-^ cells (Figure 5A), consistent with the functional observation that CD73^+^ T cells display increased longevity and increased effector function. To further identify TF networks that are involved in determining the distinct differentiation states, we compared chromatin accessibilities of CD73^+^ and CD73^-^ memory T cells by ATAC-seq. Accessibility maps differed greatly, with 3961 sites significantly more open and 2209 sites more closed in CD73^+^ cells (Figure 5B). DNA was completely inaccessible at the *NT5E* locus in CD73^-^ T cells while accessibility to CXCR5 was increased, consistent with the transcriptional data (Figure 5C).

**Figure 5:**
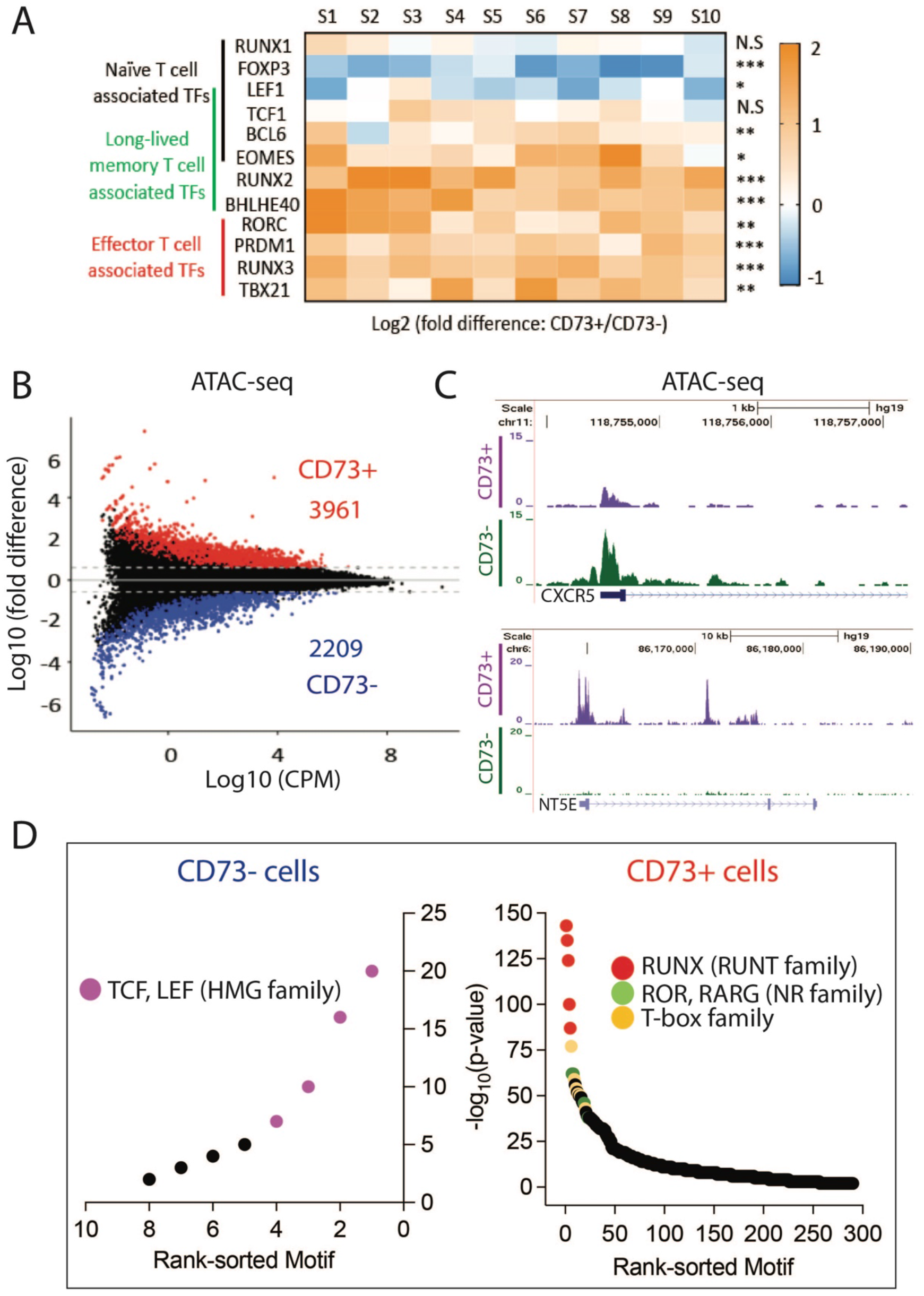
Differential chromatin accessibilities in CD73^+^ and CD73^-^ memory T cells. **(A)** Expression profile of 12 T cell-related transcriptional factors in CD73^+^ vs CD73^-^ memory CD4 T cells from 10 healthy individuals (S1 to S10) as determined by RT-PCR. Data were compared by two-tailed paired t-test. *p<0.05, **p<0.01, ***p<0.001, N.S: not significant. **(B)** Chromatin accessibility in CD73^+^ and CD73^-^ memory CD4 T cells from three healthy individuals determined by ATAC-seq. Results are shown as MA plot with differentially accessible sites (≥1.5-fold difference, p<0.05) indicated by red (more open in CD73^+^ cells) and blue (more open in CD73^-^ cells). CPM: count per million reads. **(C)** Chromatin accessibility tracks across the genes *NT5E* encoding CD73 and *CXCR5* in CD73^+^ (purple) and CD73^-^ (green) memory T cells. **(D)** TF motif enrichment at sites with decreased (left) and increased accessibility (right) in CD73^+^ vs CD73^-^ cells. Colors indicate TF families with shared motifs.

HOMER analysis of differentially accessible union peaks yielded a highly significant enrichment for RUNT (RUNX2, RUNX3), NR (RORC, RORA, RARG) and T-box family (EOMES, T-bet) TF motifs in CD73^+^ cells, while HMG family TFs (LEF1, TCF1) were the only motif enriched in CD73^-^ cells (Figure 6D). We integrated the differences in transcriptomes and in chromatin accessibilities with public ChIP-seq and T cell-specific chromatin interaction data using TF-regulatory region-target gene triplet inference modeling to construct the signature networks of key TFs and their target genes for the two T cell subsets^25,26^ (Suppl. Figure 3). The center of the major network of CD73^+^ T cells was formed by RUNX2 and RUNX3, while one smaller network was centered on EOMES, both of them indicative of CD73^+^ T cells being long-term memory T cells^10,11^ (Figure 6A). *NT5E* was the target gene with most significant inference, additional target genes included *ADAM23, MATK, CFH* and *ITGA1*. RUNX2 silencing and overexpression confirmed the role of RUNX2 in controlling NT5E transcription in CD4 as well as CD8 T cells (Figure 6B and C). RUNX3 silencing did not have an effect on CD73 expression (Figure 6D); however, forced overexpression clearly upregulated NT5E transcription (Figure 6E). Additional clusters were centered on RORC/RARG and ETS2. RORC in mature T cells is pivotal for induction and maintenance of TH17 effector T cells. The retinoic acid receptor encoded by RARG promotes the differentiation and homing of gut-resident memory CD8 T cells^27,28^. Thus, CD73 expression is a biomarker of a unique constellation of TFs in these cells. However, none of these TFs changes with age. Thus, the diminution of CD73+ T cells is not just a reflection of the regulation of CD73 expression, but is a true decline in one functional T cell subset.

**Figure 6:**
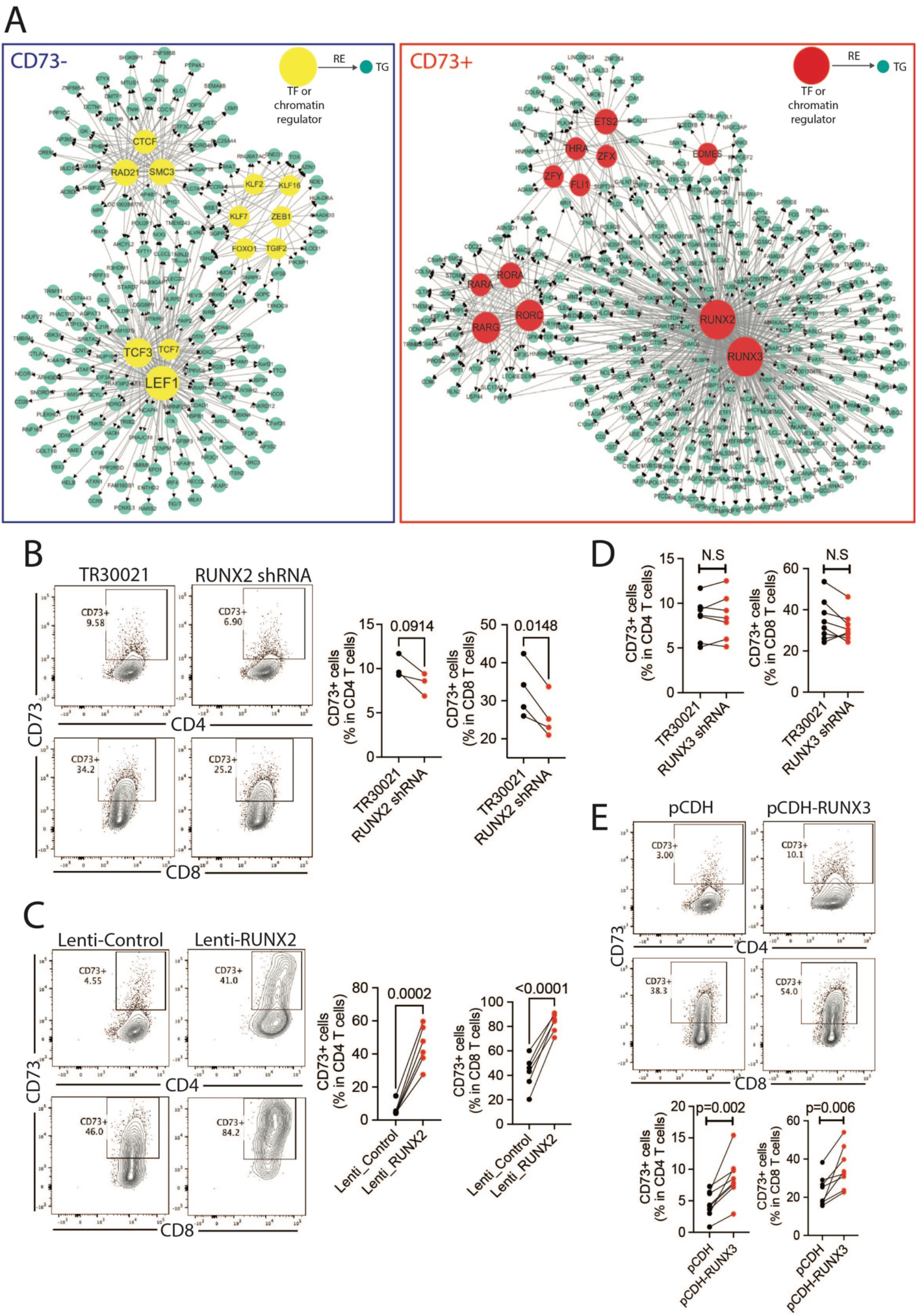
CD73^+^ and CD73^-^ memory T cells are governed by distinct transcription factor networks. **(A)** Transcription Factor-Regulatory Element-Target Gene (TF-RE-TG) networks in CD73^-^ (left) or CD73^+^ cells (right) were modeled as described in Suppl. Figure 3. Red and yellow nodes represent transcriptional factors (TF) or chromatin regulators (CR); the green nodes represent their target genes (TG) that are differentially expressed in CD73^+^ and CD73^-^ memory T cells. The size of TF nodes corresponds to the number of TF connections. (**B-D**): Freshly isolated human total T cells were activated and infected with GFP+ lentivirus containing RUNX2 shRNA (B) RUNX2 cDNA (C), RUNX3 shRNA (D), and RUNX3 cDNA (E) respectively. TR30021, pCDH and Lenti-Control served as respective controls. Transduced cells were cultured for 7 days, before CD73 expression in gated GFP^+^ cells was assessed. Results are compared by two-tailed paired t-test. N.S: not significant.

In contrast to CD73^+^ T cells, the major network in CD73^-^ T cells centered on HMG family TFs including LEF1, TCF3 and TCF7 that are known to be important for stem-like memory T cells (Figure 6A). A second smaller cluster centered around several TFs that share the functional property of being involved in dampening T cell effector functions. These genes include the transcriptional repressors TGIF2 and ZEB1, several members of the KLF family and FOXO1. These patterns are unexpected for cells that appear to be under increased turnover. A third cluster centered around CTCF together with the cohesion components RAD21 and SMC3, possibly indicating differences in chromatin structural maintenance related to the increased mitotic activity of CD73^-^ T cells.

### CD73^+^ memory T cells are prone to differentiate into cells expressing a tissue-resident memory (T_RM_) phenotype

Among TFs differentially expressed or identified in the network analysis for CD73^+^ cells, at least four were described as critical for T_RM_ cell differentiation, function or survival, including PRDM1^29^, RUNX3^30,31^, BHLHE40^14,32^ and RORC^33^. To examine whether CD73^+^ and CD73^-^ memory T cells have equal potential to differentiate into T_RM_ cells, we cultured freshly isolated human memory T cells under sequential TCR and TGFβ/IL-15 stimulation^28,34^. After differentiation, CD73^+^ cells had elevated cell surface expression of T_RM_ markers compared to CD73^-^ cells (Figure 7A/B and Suppl. Figure 4A/B). For CD4 CD73^+^ T cells, the combination of CD69 and CXCR6 was the most frequent phenotype. Expression of CD103 was less frequent; however, up to 5-10% of CD4^+^CD73^+^ cells co-expressed CD69 and CD103. For CD8^+^ CD73^+^ cells, the T_RM_-associated marker combinations CD69^+^CXCR6^+^ and CD69^+^CD103^+^ were about equally frequent. Conversely, CD73 expression was frequent on CD69^+^CXCR6^+^ cells, while infrequent on CD69^-^CXCR6^-^ cells (Suppl. Figure 4C).

**Figure 7:**
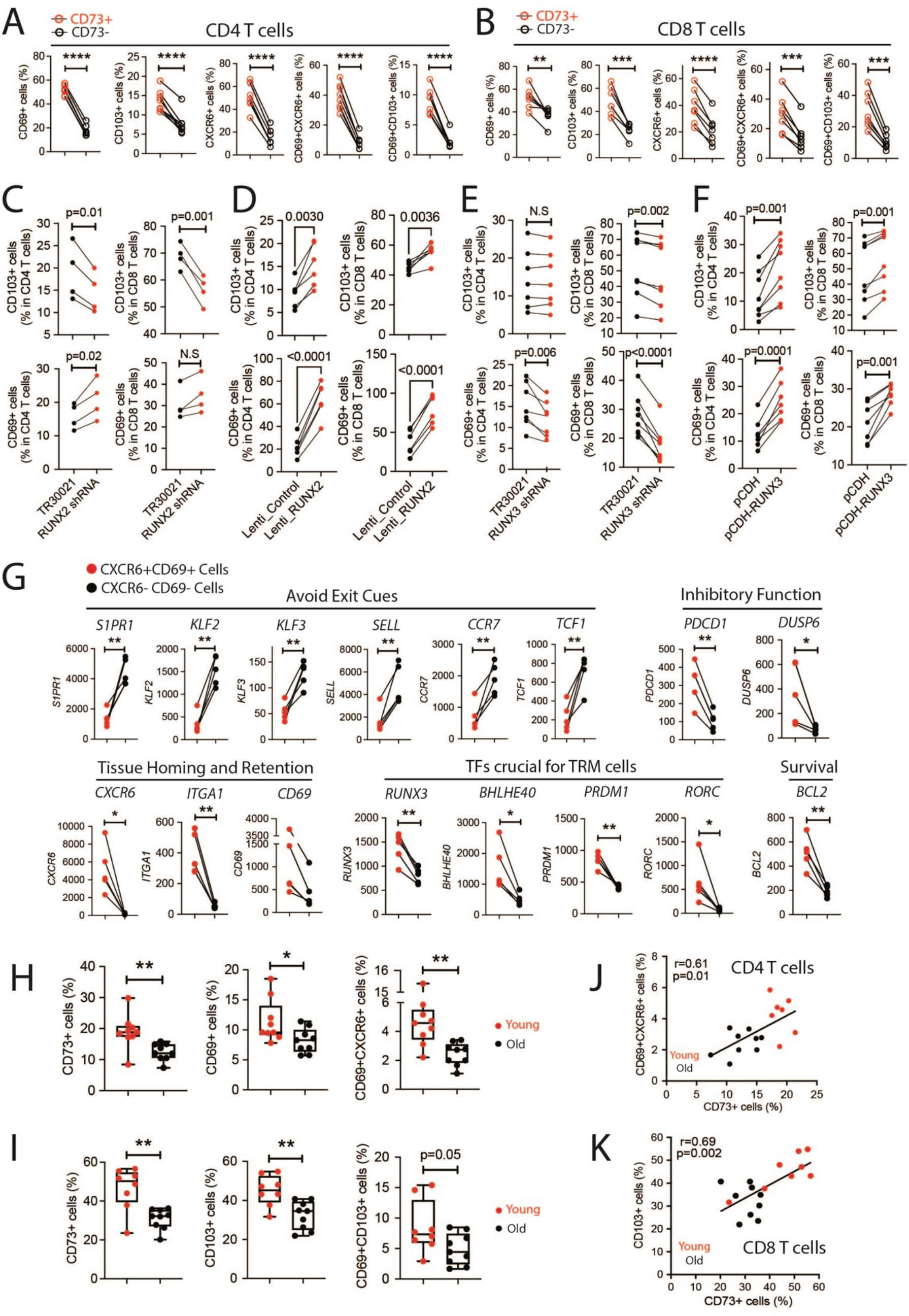
CD73^+^ memory T cells are prone to differentiate into cells with a T_RM_ phenotype. **(A and B)** Freshly isolated memory T cells were activated in vitro by anti-CD3/CD28 Dynabeads for 4 days followed by culture with TGFβ/IL-15 for 3 days. CD4 (A) and CD8 (B) T cells were analyzed by flow cytometry for the T_RM_-associated markers CD69, CXCR6 and CD103 in CD73^+^ and CD73^-^ cells. (**C-F**): Freshly isolated human total T cells were activated and infected by GFP^+^ lentivirus containing RUNX2 shRNA (C, TR30021 as a control), RUNX2 cDNA (D, Lenti-Control as a control), RUNX3 shRNA (E, TR30021 as a control) or RUNX3 cDNA (F, pCDH as a control) and differentiated under TRM development conditions for 7 days. GFP^+^ cells were gated and analyzed for CD69 and CD103 expression. **(G)** Expression profile of 16 of 19 T_RM_ core genes in the CXCR6^+^CD69^+^ and the CXCR6^-^ CD69^-^ CD4 T cell subsets that have the highest and the lowest CD73 expression, respectively. The remaining three genes (CX3CR1, S1PR5 and CRTAM) were undetectable and are not shown. qPCR results are shown as 2^(-delta Ct)^ *10^−5^. **(H-K)** Freshly isolated memory CD4 (H/J) and CD8 (I/K) T cells from young (<35y, red symbol) and older (>65y, black symbol) individuals were differentiated under 4 days of Dynabeads stimulation and 3 days of TGFβ treatment. Expression of CD73, CD69, CXCR6 and CD103 were analyzed by flow cytometry; results are summarized as box plots (H,I). Frequencies of CD73^+^ cells correlated with those of CD69^+^CXCR6^+^ cells for CD4 T cells (J) and CD103^+^ cells for CD8 T cells (K) as determined by Pearson’s correlation analysis. Data were compared by two-tailed paired or unpaired t-test. One-way ANOVA was used for multi-group comparisons. *p<0.05, **p<0.01, ***p<0.001, ****p<0.0001.

The key transcription factors regulating CD73 expression, RUNX2 and RUNX3 were also supporting the expression of TRM markers. RUNX2 silencing during TRM generation reduced the gain in CD103 expression, while keeping CD69 expression unaffected (Figure 7C and Suppl. Figure 4D). However, the forced expression of RUNX2 upregulated both CD103 and CD69 (Figure 7D and Suppl. Figure 4E) in addition to CXCR6 (Suppl. Figure 4F). RUNX3 silencing (Figure 7E and Suppl. Figure 4G) as well as overexpression (Figure 7F and Suppl. Figure 4H) documented a role for RUNX3 in CD69 and less so for CD103 expression.

We compared the two subsets differing in CD73 for the expression of 19 T_RM_ core genes^31,35,36^. Three genes were hardly detectable; all other 16 genes were consistent with the pattern described as a T_RM_ profile. *S1PR1, KLF2, KLF3, SELL, CCR7, TCF1* functioning as tissue-exiting promoters were all decreased in CXCR6^+^CD69^+^ cells (Figure 7G) that were highly enriched for CD73 positivity (Suppl. Figure 4C); *CXCR6, CD69, ITGA1* with roles in tissue homing and retention, *RUNX3, BHLHE40, PRDM1, RORC* being crucial for T_RM_ differentiation and function, *BCL2* for cell survival and the inhibitory molecules *PDCD1* and *DUSP6* were all increased in CD69^+^CXCR6^+^ cells (Figure 7G).

To control for changes in CD73 expression in the T_RM_ differentiation culture, we sorted CD73^+^ and CD73^-^ memory T cells before sequentially culturing them with TCR and then TGFβ signaling (Suppl. Figure 5). The CD73 marker was preserved before and after cell differentiation. Consistent with data shown in Figure 7, CD69, CXCR6 and CD103 T_RM_ markers were significantly elevated in CD73^+^ compared to CD73^-^ cells (Suppl. Figure 5A and B). Moreover, compared to CD73^-^ cells, the expression profile of 11-12 T_RM_ signature genes in CD73^+^ cells was consistent with T_RM_ features with upregulation of tissue homing and retention molecules (e.g., CD49a, CRTAM, CD69 and CXCR6 as well as additionally CD101 and CD103 for CD8 T cells) and downregulation of genes involved in tissue exiting (e.g., CX3CR1, KLF2, KLF3, S1PR1, S1PR5 and SELL, Suppl. Figure 5C and D).

As shown in Figure 1, the size of the CD73^+^ memory cell compartment declines with age. If T_RM_ derive from CD73^+^ T cells, the loss in CD73^+^ T cells should result in a lesser generation of T_RM_ with age. T_RM_ were generated in vitro from memory cells of adults younger than 35 years or older than 65 years as described above. After differentiation, old adults continued to have fewer CD73^+^ T cells in the CD4 as well as the CD8 T cell subset (Figure 7H/I). Generation of T_RM_ was reduced in parallel. CD69^+^CXCR6^+^ cells were around 5% for young compared to less than 3% for old memory CD4 T cells (Figure 7H). Similarly, the frequency of CD103^+^ cells was higher for memory CD8 T cells from young than from old adults (Figure 7I). Frequencies of CD73^+^ cells highly correlated with those of CD69^+^CXCR6^+^ cells in CD4 T cells and with CD103^+^ cells in CD8 T cells (Figure 7J/K). Taken together, the lower frequencies of CD73^+^ cells accounted for reduced in vitro T_RM_ generation with age.

### CD73 influences T cell survival and T_RM_ differentiation in vivo

The in vitro data with human T cells convincingly showed that CD73^+^ T cells are more prone to express T_RM_ markers. To examine whether T_RM_ in human tissues are characterized by the expression of CD73, we collected spleen, lung and jejunum tissues from three organ donors older than 65 years (Figure 8A). CD73 expression on CD4 T cells was very low in two of the three spleens (Figure 8A). Spleens from all three donors had a CD8 T cell subpopulation with high expression of CD73, two had an additional CD73^low^ population. Virtually all of the CD73^hi^ subpopulation co-expressed CD103 and CD69. Compared to splenic T cells, CD73-expressing T cells were enriched in the jejunum, especially for CD8 T cells. Up to 25% of CD45RA^-^CD4 and up to 77% of CD45RA^-^CD8 T cells expressed CD73. Virtually all jejunum T cells expressed CD73, CD73^+^ T cells frequently expressed CD103, in particular jejunal CD8 T cells, while virtually all jejunum cells expressed CD69. CD73 expression was infrequent in lung T cells, but when present, correlated with CD69 and CD103 expression.

**Figure 8:**
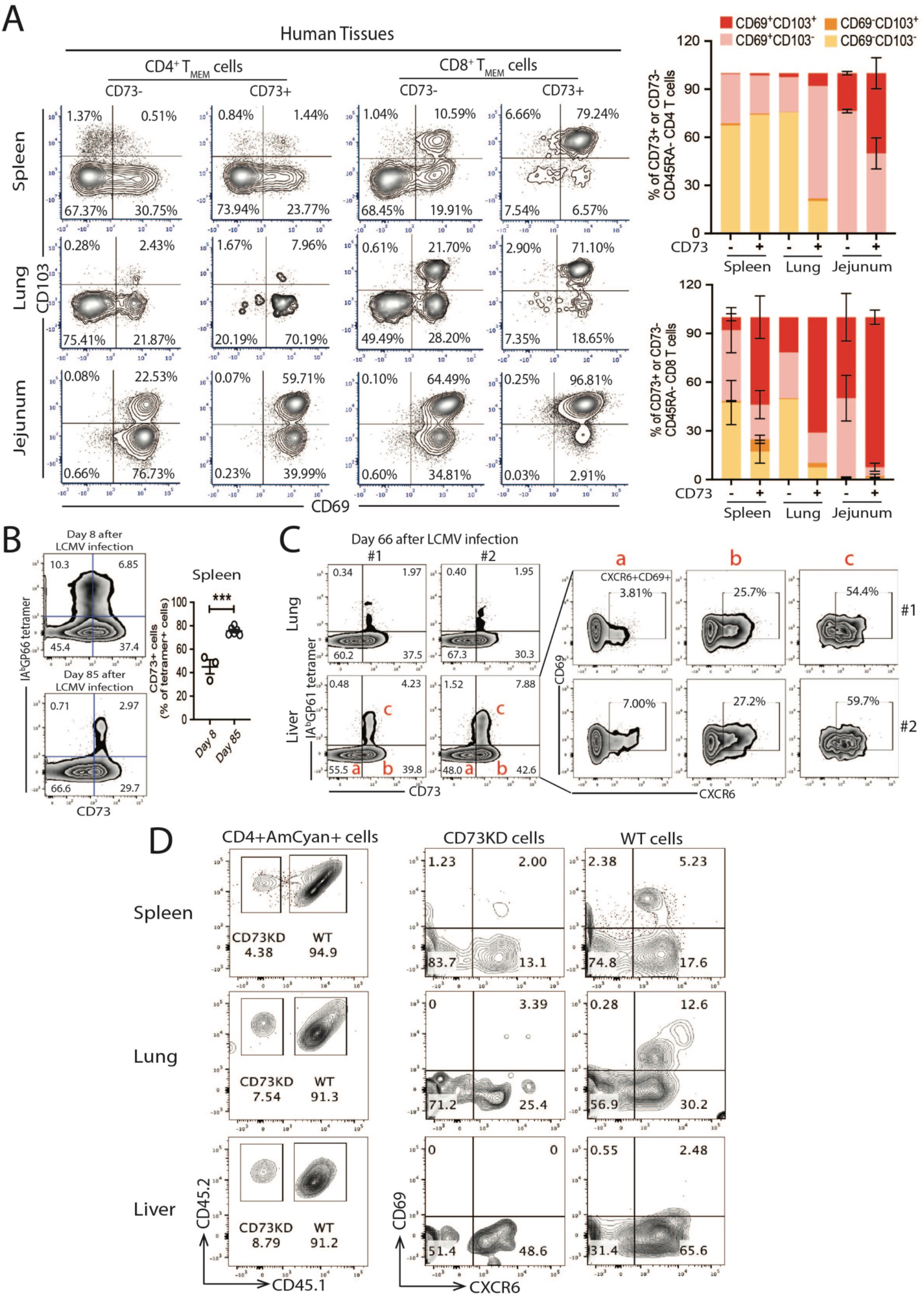
Influence of CD73 on T cell survival and T_RM_ differentiation in vivo. **(A)** Cells isolated from spleen, lung and jejunum tissues of three organ donors were analyzed for the expression of the T_RM_-markers CD69 and CD103 within CD73^+^ and CD73^-^CD45RA^-^ T cells. Representative contour plots on gated CD45^+^ cells (left panel for CD4 T_MEM_ cells; right panel for CD8 T_MEM_ cells). Subset distributions within CD73^+^ and CD73^-^ CD45RA^-^ CD4 and CD8 T cells are summarized as stacked bars. Error bars are included where data from more than one sample were available. **(B)** Mice were infected with Armstrong LCMV; CD73 expression on LCMV GP66-77 tetramer^+^ splenic CD4 T cells was determined at day 8 (effector stage) and day 85 (memory stage). Representative contour plots (left) and summary data from 3-6 mice (right). Data are compared by two-tailed unpaired t-test. ***p<0.001. **(C)** CD73 expression on tissue-resident LCMV GP61-80 tetramer^+^ CD4 T cells in lung or liver at day 66 after LCMV infection (left panel) and phenotyping of liver-resident tetramer^+^ CD4 T cells (right panel). **(D)** CD4 T cells from WT mice (CD45.2^+^) or heterozygous mice (CD45.1^+^CD45.2^+^) were infected with AmCyan^+^ retrovirus expressing CD73 shRNA or control shRNA. After sorting AmCyan^+^ cells (successfully transduced cells), congenic cells were mixed at 1:1 ratio before being transferred into WT mice through tail vein injection. After one day, mice were infected with Armstrong LCMV and sacrificed 20 days after infection before sacrifice. Transferred T cells isolated from spleen, lung and liver were analyzed for surface markers by flow cytometry. Contour plots are representative of two independent experiments.

To explore the mechanistic relationship between CD73 and tissue residency of T_RM_, we used the LCMV infection mouse model. At the effector stage on day 8 after LCMV infection, IA^b^GP66 tetramer^+^ spleen CD4 T cells varied highly in CD73 expression with about 50% of antigen-specific CD4 T cells being CD73-positive (Figure 8B). In contrast, the majority of LCMV-specific CD4 T cells at the memory stage on day 85 expressed CD73 (Figure 8B). LCMV IA^b^GP61 tetramer-specific CD4 T cells from lung and liver tissue also highly expressed CD73 (Figure 8C). Upon further phenotyping, hepatic CD73^+^ CD4^+^ T cells had a much higher percentage of CD69^+^CXCR6^+^ cells compared to CD73^-^ cells. Specifically, tetramer^+^ CD73^+^ cells co-expressed the markers characteristic for T_RM_ (Figure 8C).

To determine whether CD73 is a surrogate marker of a subset of T cells differentiating into T_RM_ or whether it is directly involved in T_RM_ differentiation and survival, we used an adoptive transfer system and LCMV infection. CD45 congenic SMARTA cells were retrovirally transduced with control- or *NT5E* shRNA. Successfully transduced cells expressing AmCyan were sorted and mixed at a ratio of 1:1 before injecting into the tail vein. Mice were then infected with LCMV. On day 20 after infection, spleen, lung and liver were analyzed for the relative percentage of each population (CD73KD vs WT cells) as well as their respective phenotypes (Figure 8D). WT cells were highly enriched compared to CD73KD cells in all three tissues, indicating improved survival. Moreover, WT cells in all three tissues expressed more CXCR6 and CD69 comparing to CD73KD cells, indicating that CD73 has direct role of CD73 in TRM differentiation (Figure 8D).

## Discussion

Here, we describe that CD73 expression distinguishes two subsets of memory T cells with markedly different epigenetic and transcriptional profiles that do not correlate with the conventional subset distinctions. CD73 is expressed on 10-40% of cells within each of the CD4 and CD8 central and effector memory T cell subsets. CD73^+^ T cells have many of the features that are associated with long-lived memory T cells including the expression of survival factors and the IL7 receptor. In parallel, they are poised to express effector molecules upon restimulation, and they differentiate into cells with the marker profile of tissue-resident memory T cells upon exposure to TGFβ/IL-15.

Memory T cells are known to not be a monolithic block but rather diverse; over the last two decades, considerable effort went into identifying functionally relevant subsets, mostly using murine infectious disease model systems. Correlating memory cell heterogeneity and the efficacy of recall responses is pivotal for understanding how protective immunity is sustained following a natural infection. Moreover, insights into the conditions that generate different memory states have the potential to find applications in vaccination strategies^5,6^.

Functional domains that are important for memory cells and that have been used to define cell heterogeneity include the homing patterns, the ability to clonally expand upon restimulation, the spectrum of effector functions and the durability. The field has heavily relied on cell surface marker profiles and here in particular, the expression of chemokine receptors. The distinction between central and effector memory cells is based on the patterns of chemokine receptors; similarly, correlation between CD4 functional lineage differentiation and chemokine receptor expression has been observed. However, recent epigenetic studies have found that the high reliance on cell surface markers may be misleading, as they only incompletely correlate with epigenetic marks such as DNA methylation or chromatin accessibility^8^.

The distinction of CD73^+^ from CD73^-^ T cells was primarily determined by a TF network that, among other genes, regulates transcription of *NT5E* encoding for CD73. We identified distinct TF - regulatory element - target gene triplet inference networks in the two subsets of memory cells using a recently described approach^26^. A core group of TFs, topped by RUNX2 and RUNX3 was characteristic for CD73^+^ T cells when compared to CD73^-^ T cells and was involved in the transcriptional regulation of *NT5E* (p<10^−53^ for enrichment). Silencing and overexpression experiments confirmed the dominant role of RUNX2 and to a lesser extent of RUNX3 in controlling *NT5E* transcription (Figure 6). Additional networks centered on EOMES that was also predicted to regulate *NT5E*, and members of the NR family, such as RARG and RORC, consistent with the notion that CD73^+^ T cells are prone to express inflammatory cytokines. A fourth cluster of target genes centered on the TFs FLI1, ETS2 and THRA, again predicted transcriptional regulators of *NT5E*, suggesting that CD73 expression is a reflection of the TF networks that control this subset of memory T cells. This does not exclude that CD73, in addition to being a biomarker of active TF networks, is important for the functional activity of CD73^+^ memory T cells.

Indeed, CD73 supported survival of antigen-specific cells after LCMV infection as well as expression of T_RM_ markers on LCMV-specific tissue-residing T cells. CD73 converts AMP to adenosine that then triggers adenosine receptors, a mechanism that has been implicated in the negative regulatory function of Treg^37^. Clearly, the population described here is not bona fide Tregs. The TF network regulating CD73 expression in T memory cells is very different from Treg cells and it is therefore not surprising that FOXP3^+^ Treg are enriched in the CD73^-^ CD4 T cell population (Suppl. Figure 1B). Whether the supporting effect on survival and T_RM_ generation observed here is related to CD73 enzyme activity and the generation of adenosine is undetermined. Different adenosine receptors exist that can transmit positive or negative signals. The main adenosine receptor expressed on T cells is the negative regulatory G protein-coupled A2AR, consistent with adenosine inducing cell inhibition^37^ or cell death in CD39 expressing short-lived effector T cells^7^. However, it is possible that dependent on the setting, A2AR stimulation can be beneficial for T cell function and survival. Stimulation of the A2AR activates PKA directing T cell differentiation^38^, protecting T cells from activation-induced cell death^39^ and preventing CD73^+^ T cells from cycling, thereby inducing quiescence^40^. Also, adenosine production in the tumor environment, generally considered immunosuppressive, has been shown to support anti-tumor responses, probably by maintaining IL7R expression and improving T cell survival^41^.

Functionally, CD73^+^ T cells displayed properties that are expected from long-lived memory cells. Their decline with age therefore could explain the defect in immune memory. CD73^+^ memory T cells have a significantly lower turnover in vivo than CD73^-^ memory T cells. In vitro, they are more resistant to undergo apoptosis, and they are more responsive to IL7 in improving survival, presumably due to increased BCL2 and IL7R expression. They exhibit increased chromatin accessibility to regulatory regions of effector genes, consistent with the previous report that memory cells display the epigenetic signature of effector T cells^42^. Upon restimulation, they are able to proliferate and efficiently produce inflammatory cytokines, frequently in combination with IL2 attesting to their polyfunctionality. Many of these features are reminiscent of stem-like memory T cells that are regulated by TCF1 cells^43^. It was therefore surprising to see that more accessible regulatory regions in CD73^+^ CD4 T cells were not enriched for TCF binding motifs; on the contrary, we found a motif enrichment of TCF1 and even more so LEF1 at sites more accessible in CD73^-^ T cells. We also saw increased expression of ribosomal genes in CD73^+^ T cells indicative of increased protein synthesis that is more characteristic of effector rather than memory cells. Subsetting memory T cells based on their CD73 expression therefore bears no relationship to the distinction between central or stem-like memory and effector memory T cells.

TF networks in CD73^+^ T cells, including the central position of RUNX3 and RORC, indicated a relationship to tissue-resident effector T cells. Indeed, CD73^+^ memory T cells were prone to differentiate in vitro upon TCR and subsequent TGFβ/IL15 stimulation into T cells expressing phenotypic markers of T_RM_. CD4^+^CD73^+^ T cells differentiated into CD69^+^CXCR6^+^ T cells, which have a gene expression profile similar to T_RM_ cells^31,35,36^; CD8^+^CD73^+^ T cells acquired the classical CD69^+^CD103^+^ T_RM_ phenotype. Thus, CD73 appears to identify a subset of circulating cells that is prone to differentiate into T_RM_ cells, possibly based on their TF networks. In vivo studies supported this interpretation. In T cells harvested from human tissue, CD103 expression was increased within the CD73^+^ T cell population, in particular for CD8 T cells from the jejunum. Studies in the murine LCMV system also support the notion that antigen-specific CD4 T cells trapped in tissue are enriched for CD73^+^ cells. In general, T_RM_ are thought to derive from short-lived effector T cells that migrate to peripheral tissue sites and stay there permanently. However, T_RM_ are also replenished from a pool of recirculating effector T cells^44-46^. Experiments in the mouse have characterized the recirculating CD8 population as intermediate positive for CX3CR1, while the corresponding CD4 population is undefined^47,48^. Our data suggest that the CD73^+^ T cell subsets of CD4 as well as CD8 T cells is highly enriched for cells able to express a T_RM_ phenotype.

Compared to CD73^+^ cells, CD73^-^ memory CD4 T cells are a more heterogeneous population, negatively defined as the absence of CD73. Transcriptome analysis suggested that they included regulatory T cells, TH2 and T follicular helper cells. Flow cytometry confirmed the enrichment for these populations, but left a large fraction of the CD73^-^ population unassigned (Figure 2 and Suppl. Figure 1C). Accordingly, TF networks did not center on lineage-specific transcription factors, but on the TCF/LEF family. However, in stark contrast to naïve and stem-like memory cells that are generally regulated by TCF/LEF, functional enrichment analysis as well as GSEA of the transcriptome provided evidence for activation and high in vivo turnover of the CD73^-^ population. Moreover, in vitro assays showed a shortened survival time, even in the presence of cytokines. These features are reminiscent of memory phenotype (T_MP_) cells that are triggered by self-antigen^49,50^ or commensal organisms. T_MP_ cells undergo a 2- to 3-fold faster turnover rate than do antigen-specific memory CD4 T cells^51-53^. Their homeostatic proliferation depends on MHC recognition and TCR signaling rather than cytokine alone^54^. Gossel et al. described a subset of memory T cells constantly replenished from naïve T cells, which may represent T_MP_^55^. Naïve T cells replaced around 10% of central memory and 1-6% of effector memory T cells every week^55^. These newly generated memory cells expressed higher level of Ki67 than previously established memory cells, indicating rapid turnover. Ki67 expression was very similar in mice from clean and dirty environments, suggesting that the proliferation was insensitive to environmental stimuli and that self-antigen recognition is the major driver for recruitment into the memory compartment^56^.

Similar to T_MP_, virtual memory (T_VM_) cells are generated independent of the recognition of exogenous antigen, however, mostly only from naive CD8 T cells. CD8 T_VM_ cells have poor TCR-induced IFNγ production comparing to true memory (T_TM_) T cells^57,58^. Quinn et al. showed that CD8 T_VM_ cells are mostly mono-functional^4^. Similarly, CD73^-^ memory T cells produce less IFNγ, and the majority of cells are mono-functional (Figure 2). Transcriptional profiling also showed elevated expression of IL-15Rβ in CD73^-^ memory CD4 T cells. IL-15 signaling is known to play a pivotal role in T_VM_ cell generation^59,60^ and high expression of IL-15Rβ (CD122) has been considered a marker for T_VM_ cells. The proportion of T_VM_ cells in the mouse increases with age as does the fraction of CD73^-^ T cells in humans as reported here.

In summary, we have identified a cell surface marker, CD73 that defines a subset of memory T cells that are long-lived, highly functional and has the ability to differentiate into tissue-residing cells. This population is declining with age. CD73 may be a useful biomarker for developing strategies for better memory cell induction as well as for enrichment of relevant cells for adoptive immunotherapy for the treatment of cancer or of selected viral infections^61-63^. The finding that CD73 is important for T cell longevity and T_RM_ differentiation holds the promise of identifying interventions that improve T cell memory, in particular in older individuals.

## Materials and Methods

### Study population and cell purification

De-identified leukocyte samples from 134 blood or platelet donors younger than 35 years or older than 65 years were purchased from Stanford University Blood bank. In addition, PBMCs were obtained from 11 healthy volunteers. 8 of these individuals were younger than 35 years old. Human tissues (spleen, lung and jejunum) were procured from three organ donors older than 65 years through collaboration with LiveOnNY, the organ procurement organization for the New York metropolitan area. The studies were approved by the Stanford University Institutional Review Board, and participants gave informed written consent. As confirmed by the Columbia University IRB, use of tissue samples obtained from brain-dead (deceased) individuals does not qualify as ‘‘human subjects’’ research.

Untouched total T or isolated CD4^+^ or CD8^+^ T cells were purified from buffy coats or whole blood with RosetteSep Human T or CD4^+^ or CD8^+^ T cell enrichment cocktails (STEMCELL Technologies, Cat#: 15061, 15062, 15063). Memory CD4^+^ or CD8^+^ T cells were either isolated from PBMCs by EasySep Human Memory CD4^+^ or CD8^+^ T cell enrichment kit (STEMCELL Technologies, Cat#: 19157 or 19159), or isolated from purified CD4^+^ or CD8^+^ T cells by CD45RO microbeads (see Suppl. Table 2). PBMCs were obtained by density gradient centrifugation using Lymphoprep (STEMCELL Technologies, Cat#: 07861). Purity of isolated cells was >90%. Human spleen, lung and intestinal samples were processed using enzymatic and mechanical dispersion to generate single-cell suspensions containing high yields of live leukocytes.

### Cell culture and flow cytometry

For surface staining, cells were incubated with fluorescence-conjugated antibodies in FACS buffer (2% FBS in PBS) for 30 min at 4°C. Cells were activated with plate-coated anti-CD3/anti-CD28 (BioLegend, OKT3 and CD28.2 clones, Cat#: 317326 and 302934) overnight before staining with anti-CD69 antibody (see Suppl. Table 2). For intracellular cytokine assays, cells were stimulated with 50 ng/ml phorbol 12-myristate 13-acetate (PMA, Peprotech, Cat#: 1652981) and 500 ng/ml ionomycin (Peprotech, Cat#: 5608212) in the presence of Brefeldin A (GolgiPlug, from BD Cytofix/Cytoperm plus kit) for 4 hours at 37°C. Cells were then sequentially incubated with surface antibody cocktail, fixed and permeabilized with Cytofix/Cytoperm plus kit (BD Biosciences, Cat#: 555028) and finally stained with fluorescence-labeled antibodies specific to the indicated cytokines. For staining of Ki67, cells were fixed with Cytofix buffer (BD Biosciences) for 10 min at 37°C, followed by permeabilization with pre-chilled Perm buffer III (BD Bioscienes) for 30 min on ice, finally stained by Ki67 antibody (see Suppl. Table 2) for 45 min at 4°C. Staining for FOXP3 followed the instruction provided by True-Nuclear Transcription Factor Buffer Set kit (BioLegend, Cat#: 424401). For apoptosis analysis, cells were incubated for the indicated time in the absence or presence of IL-7 (10 ng/ml, Peprotech, Cat#: 200-07) or stem cell factor (20 ng/ml, STEMCELL technologies, Cat#: 78062.1), then stained with fluorescence labeled Annexin V and 7-AAD (BD apoptosis detection kit, Cat#: 559763). For in vitro T_RM_ cell differentiation, purified memory T cells were stimulated with anti-CD3/CD28 Dynabeads (ThermoFisher Scientific, Cat#: 11132D; bead to cell ratio 1:2) for 4 days, followed by TGFβ alone or together with IL-15 (TGFβ: 10 ng/ml, STEMCELL Technologies, Cat#: 78067; IL-15: 10 ng/ml, Peprotech, Cat#: 200-15) for 3 more days. Cells were stained with antibody cocktail specific to CXCR6, CD69, CD103 (see Suppl. Table 2). For flow-cytometric analysis of cells from human tissues, single-cell suspensions were pre-incubated with Fc Block (BioLegend, Cat#: 422302), stained with fluorochrome-conjugated antibodies and fixed with FOXP3/Transcription Factor Fixation Buffer (Tonbo Biosciences, Cat#: TNB-1020-L050). Tissue-derived cells were gated to exclude dead cells, gate on singlets, followed by gating on CD45^+^, CD3^+^, and finally CD4^+^ or CD8^+^ T cells to assess CD45RA, CD73, CD69 and CD103 expression. Single cell suspensions from mouse spleen, lung and liver were stained with IAb LCMV GP66-77 DIYKGVYQFKSV (GP66) and IAb LCMV GP61-80 GLKGPDIYKGVYQFKSVEFD (GP61) tetramers, obtained from the NIH tetramer core facility (Emory University, Atlanta, GA), followed by staining with antibodies for cell surface markers (see Suppl. Table 2). Dead cells were excluded from the analysis using live/dead fixable aqua (Invitrogen, Cat#: L34966). In adoptive cell transfer experiments, T cells were isolated from mouse lung and spleen and then incubated with antibody cocktail specific to CD45.1, CD45.2, CXCR6, CD69 for surface staining (see Suppl. Table 2).

Cells were analyzed on an LSRII or LSR Fortessa (BD Biosciences); flow cytometry data were analyzed using FlowJo (TreeStar) or FCS Express (De Novo Software). Cell sorting was done on a FACS Aria2 or Aria3 (BD Biosciences).

### Lentiviral transduction of human T cells

RUNX3 cDNA was generating using the primers set hRUNX3_enzy_F: 5’ AACTAGCTAGCatggcatcgaacagcatcttcg 3’ and hRUNX3_enzy_R: 5’ ATACGCGGATCCtcagtagggccgccacac 3’ and cloned into a commercial lentivector containing a GFP reporter gene (pCDH-GFP-Em-CD513B-1 from System Biosciences). To overexpress RUNX2 in human T cells, we purchased commercial RUNX2/GFP lentivector (Origene, Cat#: RC212884L4). To knockdown RUNX3 or RUNX2 in human T cells, we used RUNX3 or RUNX2 human shRNA plasmid containing GFP reporter gene (Origene, Cat#: TL309682 for RUNX3; Cat#: TL309683 for RUNX2). Lentivirus was produced by transfection of the lentiviral vector, along with psPAX2 (Plasmid #12260; Addgene) and pMD2.G (Plasmid #12259; Addgene) expression vectors into HEK293T cells using Lipofectamine LTX (ThermoFisher Scientific). Lentiviral particles were collected 48 and 72 hours after transfection, filtered through a 0.45 um syringe filter (Millipore), concentrated using Peg-it solution (System Biosciences) and titrated on HEK293T cells. For lentiviral transduction, T cells were activated with anti-CD3/anti-CD28 beads and cultured with a lentiviral vector expressing scrambled control or target plasmids, at a multiplicity of infection of 10 in the presence of 8 mg/ml polybrene (Sigma) and 10 U/ml human IL-2 (Peprotech). After 48 hours, medium was changed once and cells cultured for a total 4 days. Cells were then cultured in medium containing IL-15 and TGFβ for another 3 days for TRM induction.

### Murine LCMV model

C57BL/6J (B6) mice were purchased from the Jackson Laboratory. LCMV-Armstrong was grown in BHK cells and titered in Vero cells. Male mice, 8-10 weeks of age, were adoptively reconstituted with transduced SMARTA cells and infected i.p. with a dose of 2 × 10^5^ or 5× 10^5^ plaque-forming units (PFU). Retroviral transduction was performed as follows: *Nt5e* shRNAmir (CAGGTTGAGTTTGATGATAAAG) was inserted into pLMPd-Amt vector. Virions were packaged in the Plat-E cell line; the medium was replaced after 10 hours, retroviral supernatant was collected after 48 hours. CD4^+^ T cells were purified from CD45.1^+^ CD45.2^+^ or CD45.2^+^ SMARTA splenocytes by negative selection (STEMCELL Technologies) and stimulated in 12-well plates pre-coated with 8 μg/mL anti-CD3 (145-2C11; eBioscience) and 8 μg/mL anti-CD28 (37.51; eBioscience) antibodies. After 24 hours, cells were transduced with supernatant containing retrovirus (CD45.1^+^ CD45.2^+^ SMARTA with control virus and CD45.2 SMARTA with *Nt5e* shRNAmir virus) in the presence of 8 μg/mL polybrene (MilliporeSigma) by centrifugation for 90 min at 1500 g at 32°C. Twenty-four hours after transduction, AmCyan-positive cells (successfully transduced cells) were sorted, SMARTA cells transduced with the *Nt5e* shRNAmir and control shRNAmir retroviruses were mixed at a 1:1 ratio and a total of 1×10^5^ cells were injected into recipient mice through tail vein. After resting for 1 day, recipient mice were infected with 2×10^5^ PFU LCMV Armstrong. Mice were sacrificed for cell analysis in spleen, lung and liver 20 days after infection. Lung were digested by collagenase type I (Sigma, Cat#: SCR103) for 1 hour at 37 degree, then crushed and filtered to obtain a single cell suspension. Liver and spleen were directly crushed and filtered into single cells without the enzyme digestion step. Cells were pelleted at a speed of 2000 rpm for 5 min and suspended in 44% Percoll and layered on the top of 67% Percoll. After centrifugation at the speed of 2200 rpm for 22 min, T cell layers were harvested and washed several times before surface marker staining and flow cytometric analysis. All mice were housed in the Stanford Research Animal Facility according to Stanford University guidelines. Animal experiments were approved by the Stanford University Institutional Animal Care and Use Committee.

### RNA isolation and quantitative RT-PCR

Total RNA was isolated using either RNeasy Plus Mini or Micro kit (QIAGEN, Cat#: 74134 or 74034), depending on the cell number, and converted to cDNA using High-Capacity cDNA Reverse Transcription Kit (ThermoFisher Scientific, Cat#: 4368813). Quantitative RT-PCR was performed on an Eppendorf Thermal Cycler using Powerup SYBR Green Master Mix (ThermoFisher Scientific, Cat#: A25776) according to the manufacturer’s instructions. Expression levels were normalized to *ACTB* expression and displayed as 2^-ΔCt^ *10^−5^. Primer sequences are shown in Suppl. Table 1.

### Lymphocyte isolation from human tissues

Tissue samples were maintained in cold saline and brought to the laboratory within 2-4 hours of organ procurement as described^64,65^. Spleen samples were chopped up, incubated in enzymatic digest solution (RPMI medium containing 10% FBS, L-glutamate, sodium pyruvate, nonessential amino acids, penicillin-streptomycin, collagenase D [1 mg/ml], trypsin inhibitor [1 mg/ml], and DNase I [50-100 μg/ml]) for 1.5 hours, then mechanically disrupted using a tissue homogenizer (Bullet Blender), filtered through a tissue sieve, and enriched for mononuclear cells using Ficoll density gradient centrifugation. Lung samples were processed as above except for the use of a tissue dissociator (gentleMACS) instead of a homogenizer to mechanically disrupt the samples. Jejunum was separated from intestinal samples after removal of mesenteric lymph nodes. After cleaning off fatty tissue, intestinal tissue segments were washed with PBS and injected with enzymatic digest solution. The segments were then chopped into small pieces and incubated in enzymatic digest solution. After 1.5 hours of incubation, tissue digests were mechanically disrupted using a tissue dissociator (gentleMACS), filtered through a tissue sieve, and enriched for mononuclear cells by density gradient centrifugation using 40% Percoll. The resulting cell suspensions containing high yields of live leukocytes were resuspended in complete RPMI medium.

### RNA-seq and data processing

RNA was extracted using RNeasy Plus (Qiagen, Cat#: 74034) from 300,000 to 500,000 sorted CD73^-^ and CD73^+^CD45RO^+^CD4^+^T cells. Ribosomal RNA was removed from each RNA extraction using Ribo-Zero Gold rRNA Removal kit (Illumina, Cat#: MRZG12324). RNA-seq libraries were generated by TruSeq Stranded mRNA Library Prep kit (Illumina, Cat#: 20020594) and sequenced on an Illumina HiSeq 4000 sequencer. Sequencing reads were mapped to human genome hg19 using STAR^66^. RPKM values were called using HOMER (http://homer.salk.edu/homer/motif/) analyzeRepeats.pl program. Differential expression was performed with DEseq2 using the raw counts of genes associated with each sample generated from HOMER.

### Gene set enrichment analysis (GSEA)

Gene expression data from the RNA-seq analysis of CD73^-^ and CD73^+^CD45RO^+^CD4^+^T cells were compared to a priori defined gene sets following standard protocols (http://www.broad.mit.edu/gsea/).

### ATAC-Seq library preparation, sequencing, and data preprocessing

50,000 sorted CD73^-^ and CD73^+^CD45RO^+^CD4^+^T cells were subjected to Omni-ATAC^67^ to profile the accessible chromatin landscape. ATAC-seq libraries were sequenced on Illumina HiSeq4000 sequencers. ATAC-Seq pair-end reads were trimmed off Illumina adapter sequences and transposase sequences using a customized script and mapped to hg19 using bowtie^68^ with parameters –S –X2000 –ml. Duplicate reads were discarded with samtools rmdup^69^. Peaks were identified using MACS2 with -f bed -q 0.01 –nomodel --shift 0. Overlapped peaks from all samples were merged into a unique peak list, and raw read counts mapped to each peak for each individual sample were quantified. Differentially accessible peaks from the merged union peak list were identified with the edgeR package (Bioconductor) using raw counts of each samples in the union peak list with a fold change threshold of 1.5, and a p-value <0.05.

### Transcription factor motif enrichment analysis of differentially accessible sites

Transcriptional factor motifs enriched for selected peak compared to background regions were identified using HOMER “findMotifsGenome.pl” using default parameters (http://homer.salk.edu/homer/motif/). TF motifs were ranked based on the -log_10_ (p-value) of the enrichment level.

### Modeling of Transcription Factor-Regulatory Element-Target Gene (TF-RE-TG) networks

We used a previously described inference model of TF-RE-TG regulatory networks^26^ by integrating RNA-seq, ATAC-seq and human CD4 T cell-specific Trac-looping data^25^ to assess the network differences in the two memory cell populations. The model quantifies the interaction of each RE with relevant TFs to affect the expression of their TG. We started with an assembled union peak list called from ATAC-seq across all samples as putative REs, identified the upstream TFs and downstream genes for a RE, treated each TF-RE-TG triplet as the basic regulatory unit, ranked them by integrating genomic features, and extracted the significant regulatory relations. RE openness was defined as the fold enrichment of the read starts in this region versus the read starts in a 1M bp background window. Each TF was described by its motif binding score to the RE and its expression level from the RPKM value of RNA-seq. The relationship of a TG with a RE were derived from the physical interactions measured by the loops called from Trac-looping data^25^. The computation involves four major steps.

### Step 1: Finding TF-RE-TG triplet

For TF-RE pairs, we used HOMER to scan differentially accessible regions to find all positions with substantial similarity to TF’s sequence motif or position weight matrix (PWM) and assemble TF-RE pairs. For RE-TG pairs, we considered both proximal and distal regulation. In each population, we used HOMER to annotate differential peaks with nearest genes and select peaks within 5Kb of TSS to construct the proximal RE-TG regulation pair. As for distal regulation, we introduced Trac-looping data in CD4^+^ T cells^25^, selected region-region pairs if either region was located in promoter, and expanded both regions in the pair by 2.5kb into both directions. Then, we overlapped the differential peaks with those selected pairs of Trac-looping data to construct the distal RE-TG regulation pair. We aggregated the two sets of pairs together as the predicted RE-TG sets. Through matching TF-RE pair and RE-TG pair, we constructed all candidate TF-RE-TG triplets in the network.

### Step 2: Collecting genomic features from ATAC-seq and RNA-seq data

After identifying all TF-RE-TG triplets, we determined a regulatory score and ranked triplets for both populations. Scores were derived from the following variables in the RNA-seq and ATAC-seq data: differences in TF and TG expression, differences in openness of REs, TF binding derived from motif occurrence.

#### 1. Expression of TF and TG

Gene expression was quantified as RPKM (Reads Per Kilobase of transcript per Million reads mapped). Fold change of total number of reads mapped to TF or TG (M_i_) with gene length L_TF_ or L_TG_ to the totally mapped reads N in the experiment,

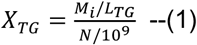

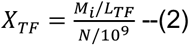

Superscript i,j were added to distinguish the expression of TF and TG in sample i and j(X^i^_TF,_X^j^_TF_). Thus, we obtained the gene expression and calculated its fold change as max {1, average B} / max {1, average A}, where average B and average A are the average expression values in B and A populations. Fold difference over 1 means increase, fold difference below 1 means decrease, and fold differences=1 means no obvious expression difference (i.e. RPKM<=1) detected among all samples.

#### 2. REs’ openness

The RE is more likely to be in the open chromatin region if TF utilizes this RE to regulate TG. We defined openness for RE from ATAC-seq. ATAC-seq measures the count of reads in a given region. We quantified the openness for the RE e_i_ by a fold change score, which computed the enrichment of read counts in e_i_ by comparing with a larger background region. Briefly, let N_i_ be the number of reads in RE e_i_ of length L_i_ and G_i_ that in the W background window around this RE. The openness of RE e_i_ can be defined as

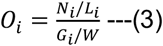

#### 3. Motif enrichment score

We used HOMER for motif enrichment analysis on differentially accessible peaks.

We set the -log(p-value) of each motif from homer outputs as enrichment score E^i^_TF_ which represents the enrichment of TF’s corresponding motif in differentially accessible peaks in sample i.

#### 4. TF activity score

We defined TF’s activity score TFA_i_ to represent the activity of TF in sample i, which combines its expression change and corresponding motif’s enrichment score. X^r^ is the TF expression level in reference. It is formally defined as:

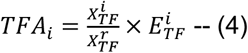

### Step 3: Integrate genomic features and rank TF-RE-TG triples

Our aim was to model how a TF regulated a TG via REs with conditions measurement in matched ATAC-seq and RNA-seq data. For a given TF-RE-TG triplet, we assumed that TF regulated this TG’s expression by REs. Thus, we can collect the genomic features for TF-RE-TG triplets in CD73^+^ and CD73^-^ cells and calculate the fold change for openness of RE and expression of TF and TG.

With those features, we assumed a normal distribution for each feature across all triplets and independence of transformed features. Using Fisher’s method, we combined the features into score S.

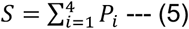

Where P_i_ is the -log(p-value) for the i-th hypothesis to assess the significance level of feature i. When the p-value tends to be small, the test statistic S will be large, which suggests that TF-RE-TG regulation is significant. S follows a chi-squared distribution with 2K degrees of freedom, from which a p-value for the global hypothesis can be easily obtained, K is the number of features being combined (K=4 in our case). As a result, all the triplets can be ranked by score S and score S converted into p-value.

### Step 4: Extracting significant TF-RE-TG triplets into network for visualization

By taking a cutoff p-value<0.05, we predicted a set of TF-RE-TG triplets. Pooling all the triplets together, we then have a TF-RE-TG network, where TF and TG are nodes, and RE is the edge. We counted the node number for TF and ranked them in each population. TF-TG networks were visualized by Cytoscape^70^.

### Statistical analysis

Statistical analysis was performed using Prism (GraphPad). Paired or unpaired two-tailed Student’s t-tests were used for comparing two groups. A two-tailed Pearson’s correlation test was used for correlation analysis. One-way ANOVA with Tukey’s post hoc test was used for multi-group comparisons. p<0.05 was considered statistically significant.

## Acknowledgements

This work was supported by the National Institutes of Health (R01 AR042527, R01 HL117913, R01 AI108906, R01 HL142068, and P01 HL129941 to C.M.W, P01 AI106697 to D.L.F and R01 AI108891, R01 AG045779, U19 AI057266, and R01 AI129191 to J.J.G) and with resources and the use of facilities at the Palo Alto Veterans Administration Healthcare System. Network modelling was supported by National Natural Science Foundation of China (NSFC) grants (No.11871463, No.61671444 and No.61621003 to Y.W.). N.L. was supported by NSF GRFP. The content is solely the responsibility of the authors and does not necessarily represent the official views of the National Institutes of Health. Flow cytometry and fluorescence activated cell sorting was performed in the Palo Alto Veteran Administration Flow Cytometry Core supported by the US Department of Veterans Affairs and the Palo Alto Veterans Institute for Research. Technical assistance was provided by Dr. Brandon Carter. Correlation analysis of transcriptomes between CD73^+^ and CD73^-^ cells vs T_EM_ and T_CM_ cells, were done by Feng Zhang from Shanghai Jiaotong University School of Medicine (Shanghai, China).

## Author Contributions

F.F., W.C., D.L.F., C.M.W. and J.J.G. designed research and analyzed data. F.F., W.C., N. L., L.L. and C.K. performed the experimental work. W.Z. and Y.W. performed the network analysis. S.G. analyzed high-throughput data. S.L., H.Z. and B.H. recruited donors. F.F., Y.W. and J.J.G. wrote the manuscript.

## Data Availability Statement

RNA-seq data of CD4 and CD8 T cell subsets from young and old adults are available in SRA with accession code PRJNA638216. RNA- and ATAC-sequence data from CD73^+^ and CD73^-^ CD4 memory T cells were deposited in GEO under accession number GSE157164.

