## Supplement for "Influence of age on functional memory T cell diversity"

### Supplementary Figure 1

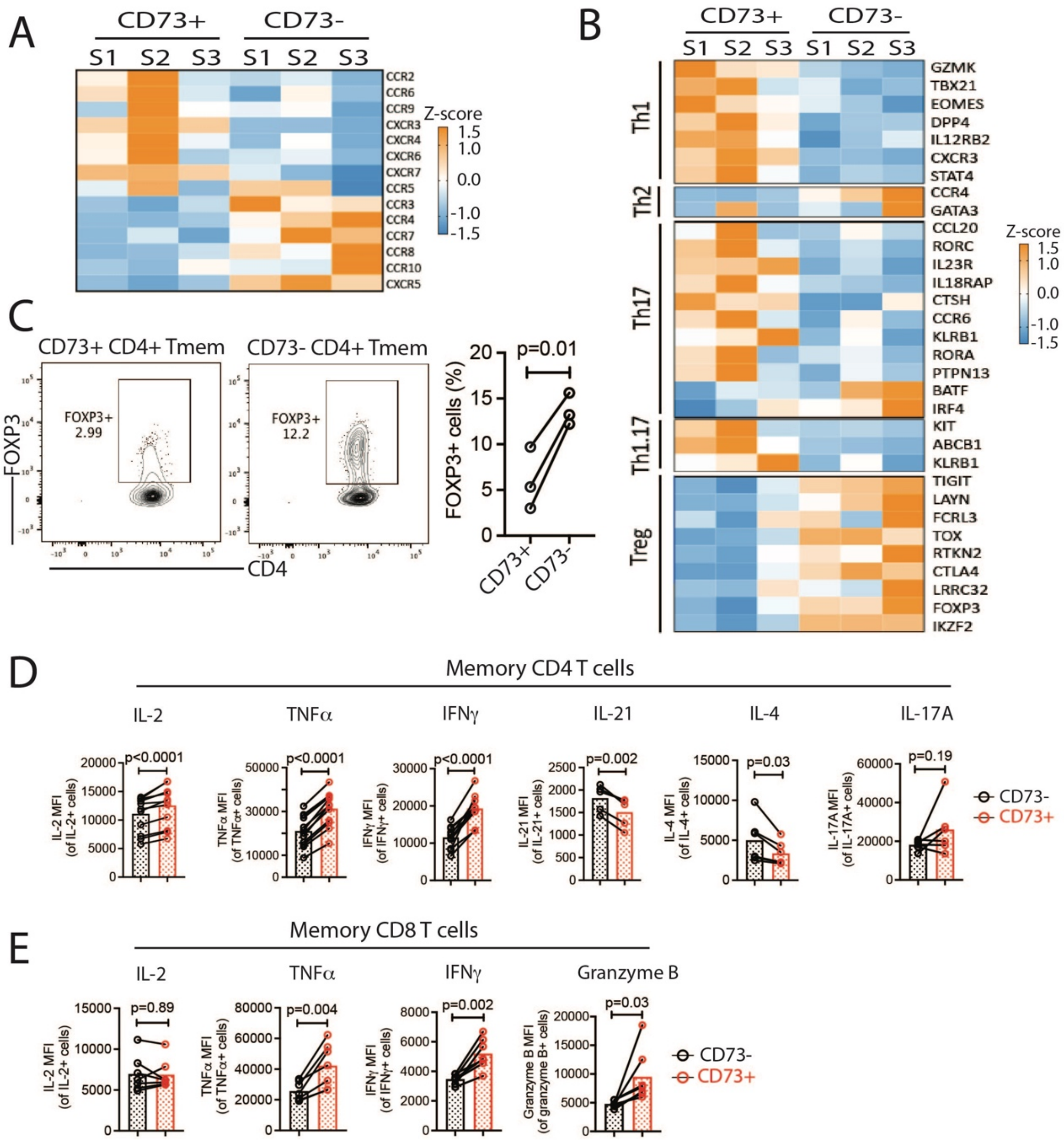


**Supplementary Figure 1, related to Figure 1C and Figure 2. Effector functions of CD73⁻ and CD73⁺ memory T cells. (A)** Heatmap of chemokine receptor gene expression in $CD73^+$ vs $CD73^-$ T cells. **(B)** Heatmap of the expression of CD4 subset-defining genes in $CD73^+$ vs $CD73^-$ cells. **(C)** Frequencies of $FOXP3^+$ cells in $CD73^+$ vs $CD73^-$ memory CD4 T cells. Representative contour plots (left) and frequencies from three experiments (right). **(D and E)** Memory CD4 (D) or CD8 (E) T cells were stimulated as described in Figures 2D and E. The expression level (MFI; median fluorescence intensity) of each cytokine in gated cytokine-positive cells was determined by flow cytometry. Data were compared by two-tailed paired t-test.

### Supplementary Figure 2

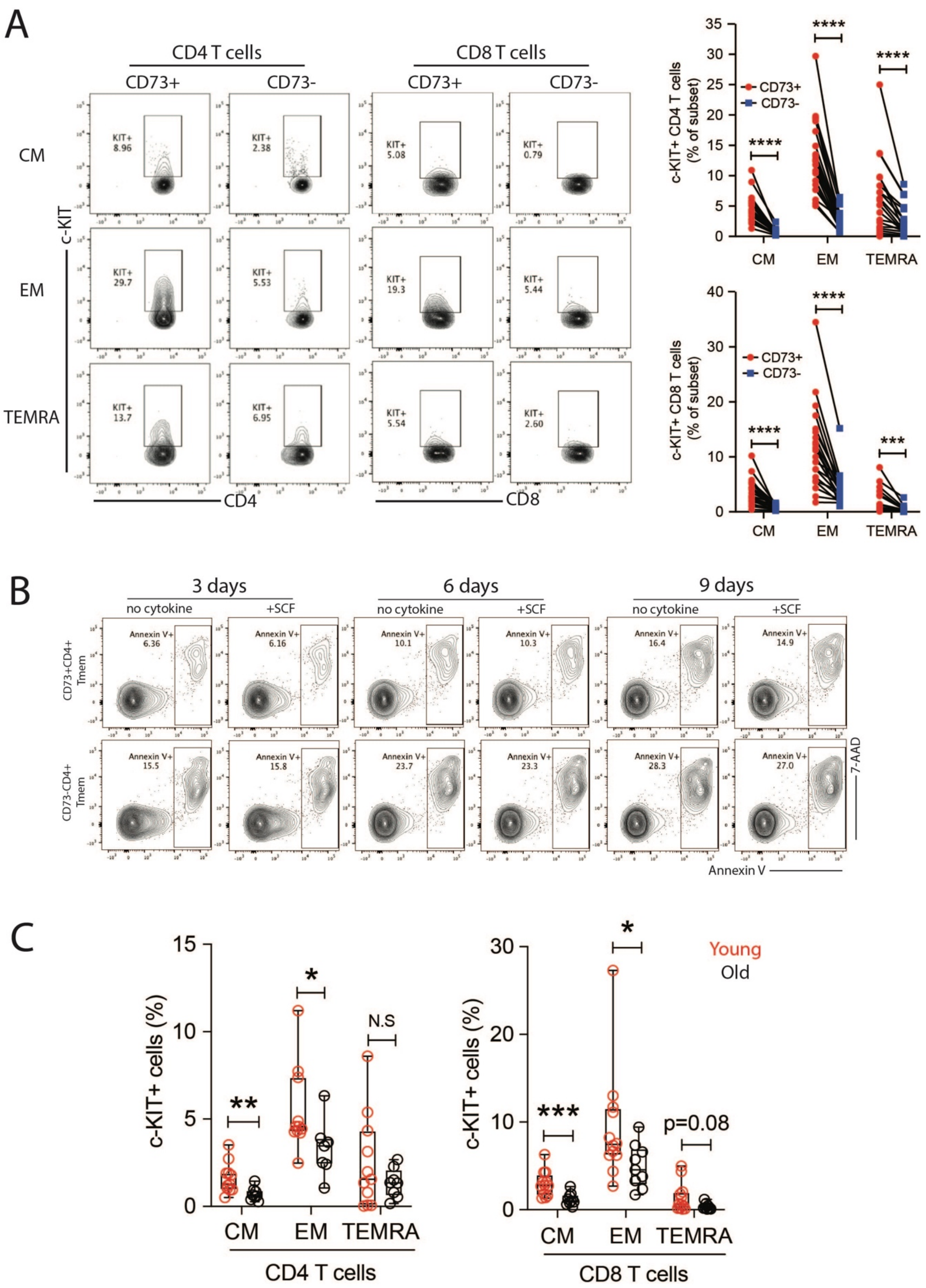

**Supplementary Figure 2, related to Figure 3: Growth factor receptor expression on $CD73^-$ and $CD73^+$ memory T cells. (A)** c-KIT expression in $CD73^+$ and $CD73^-$ CD4 and CD8 memory T cell subsets. Representative contour plots (left) and summary data (right). **(B)** $CD73^+$ and $CD73^-$ memory CD4 T cells, respectively were cultured in the absence or presence of SCF (20ng/ml). At indicated time points, cells were stained with Annexin V and 7-AAD. Contour plots representative of 3-4 independent experiments. **(C)** c-KIT expression in CD4 (left) and CD8 (right) memory T cell subsets from young (<35y) and older (>65y) individuals. Data were compared by two-tailed paired (A) or unpaired t-test (C). $*p<0.05$, $**p<0.01$, $***p<0.001$, $****p<0.0001$. N.S: not significant.

### Supplementary Figure 3

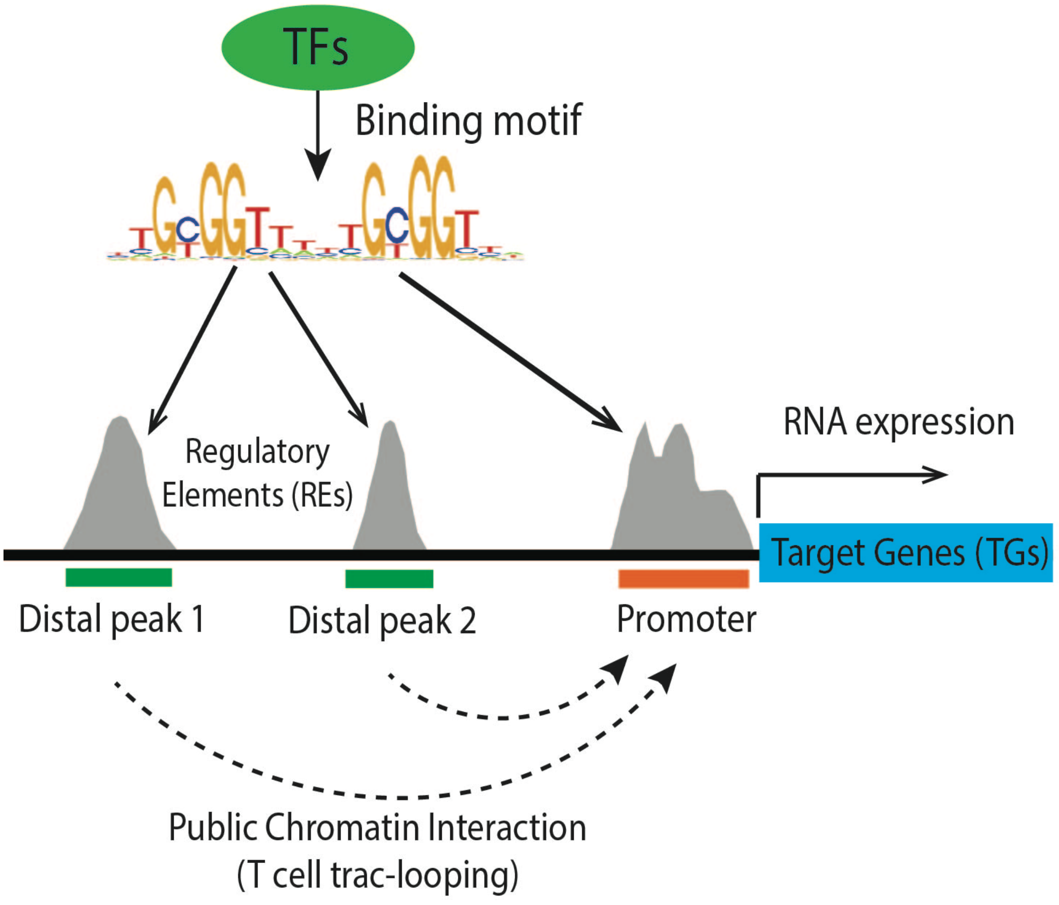


**Supplementary Figure 3, related to Figure 6: Diagram illustrating the prediction of TF-regulatory element-target gene triplet inference networks.**

### Supplementary Figure 4

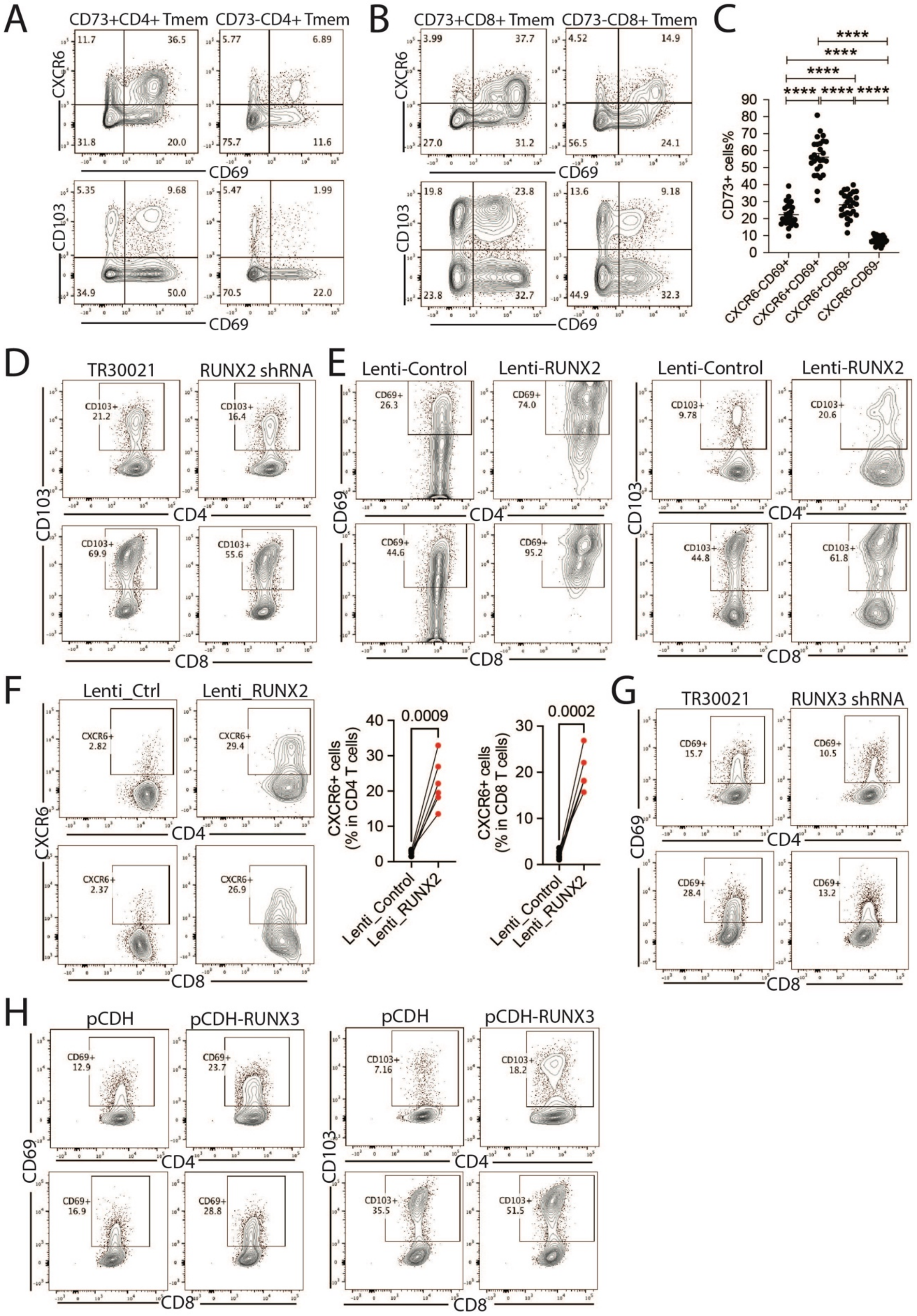

**Supplementary Figure 4, related to Figure 7A-E: Transcription factors RUNX2 and RUNX3 control expression of functional cell surface markers in $T_{RM}$ cells.**

**(A-B)** Representative contour plots for Figure 7A-B, showing the expression of CD69, CXCR6 and CD103 on $CD73^+$ and $CD73^-$ cells after $T_{RM}$ differentiation. **(C)** CD73 expression in CD4 T cell subsets defined by CXCR6 and CD69 expression level. **(D-H)** Representative contour plots for Figures 7C-F, showing CD69 and CD103 expression after RUNX2 or RUNX3 knockdown or overexpression (**D, E, G, H**). CXCR6 upregulation after RUNX2 overexpression (**F**). Representative contour plots (left) and summary data (right). Cells were analyzed on GFP+ cells.

### Supplementary Figure 5

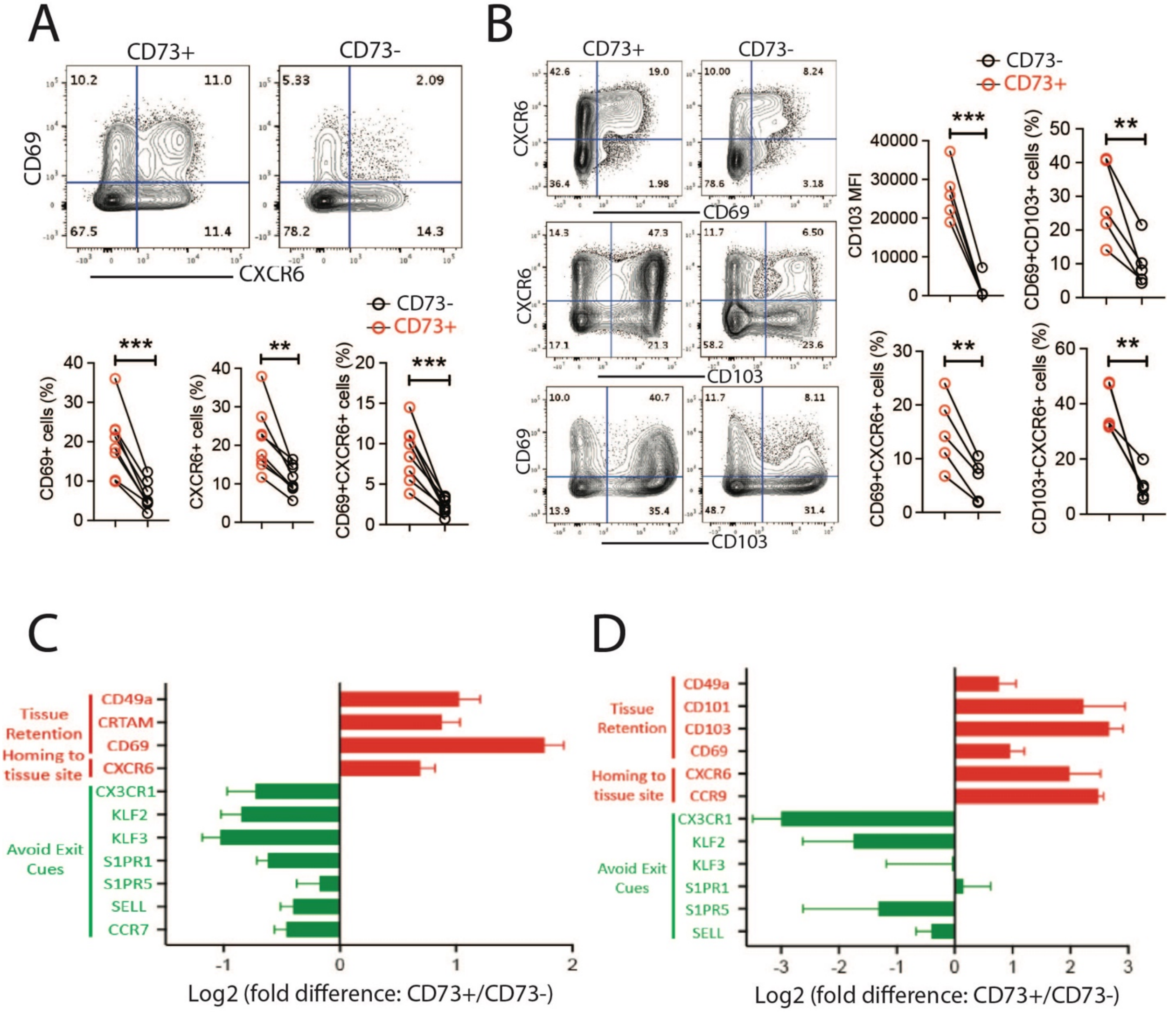


**Supplementary Figure 5, related to Figure 7A-B: Preferential differentiation of CD73$^{+}$ memory T cells into T cells characterized by a $T_{RM}$ phenotype.**

**(A-D)** Purified CD73$^{+}$ and CD73$^{-}$ memory CD4 (A/C) and CD8 (B/D) T cells were cultured sequentially with anti-CD3/CD28 Dynabeads for 4 days and TGFβ for another 3 days. CD69, CXCR6 and CD103 were analyzed by flow cytometry (A/B). Expression profile of $T_{RM}$ signature genes were determined by qPCR of CD4 (C) and CD8 (D) CD73$^{+}$ and CD73$^{-}$ cells. Data were compared by two-tailed paired t-test. **p<0.01, ***p<0.001.

**Supplementary Table 1: Primer sequences for real-time qPCR**

| Gene Name | Forward primer (5'-3') | Reverse primer (5'-3') |
|---|---|---|
| NT5E | CCAGTACCAGGGCACTATCTG | TGGCTCGATCAGTCCTTCCA |
| ACTB | ATGGCCACGGCTGCTTCCAGC | CATGGTGGTGCCGCCAGACAG |
| SELL | ACCCAGAGGGACTTATGGAAC | GCAGAATCTTCTAGCCCTTTGC |
| CCR7 | ATTTGTTTCGTGGGCCTACTG | TCATGGTCTTGAGCCTCTTGA |
| CX3CR1 | ACTTTGAGTACGATGATTTGGCT | GGTAAATGTCGGTGACACTCTT |
| ITGA1 | GCTCCTCACTGTTGTTCTACG | CGGGCCGCTGAAAGTCATT |
| CRTAM | GACGCTCACTCTAAAGTGTGTC | CTTGCAGGGTTACGTTAGGCA |
| CXCR6 | GACTATGGGTTCAGCAGTTTCA | GGCTCTGCAACTTATGGTAGAAG |
| CD69 | ATTGTCCAGGCCAATACACATT | CCTCTCTACCTGCGTATCGTTTT |
| CD101 | AAGGTTACCCAGTCAGCATTG | CTTGGTGCTAATGATCTGGACTT |
| CD103 | TGCTGGCCGCTTTCAATGT | ACAGGATGGCAAAGGATTTCAT |
| S1PR1 | TTCCACCGACCCATGTACTAT | GCGAGGAGACTGAACACGG |
| S1PR5 | GCGCACCTGTCCTGTACTC | GTTGGTGAGCGTGTAGATGATG |
| KLF2 | CTACACCAAGAGTTCGCATCTG | CCGTGTGCTTTCGGTAGTG |
| KLF3 | TGTCTCAGTGTCATACCCATCT | CCTTCTGGGGTCTGAAAGAACTT |
| PDCD1 | CCAGGATGGTTCTTAGACTCCC | TTTAGCACGAAGCTCTCCGAT |
| DUSP6 | GAAATGGCGATCAGCAAGACG | CGACGACTCGTATAGCTCCTG |
| MKI67 | ACGCCTGGTTACTATCAAAAGG | CAGACCCATTTACTTGTGTTGGA |
| BCL2 | GGTGGGGTCATGTGTGTGG | CGGTTCAGGTACTCAGTCATCC |
| IL7R | CCCTCGTGGAGGTAAAGTGC | CCTTCCCGATAGACGACACTC |
| KIT | CGTTCTGCTCCTACTGCTTCG | CCCACGCGGACTATTAAGTCT |
| RUNX3 | GCGAGGGAAGAGTTTCACCC | TTGATGGCTCGGTGGTAGGT |
| EOMES | CTGCCCACTACAATGTGTTCG | GCGCCTTTGTTATTGGTGAGTTT |
| TBX21 | GTCCAACAATGTGACCCAGAT | ACCTCAACGATATGCAGCCG |
| RUNX1 | CTGCCCATCGCTTTCAAGGT | GCCGAGTAGTTTTCATCATTGCC |
| BHLHE40 | GACGGGGAATAAAGCGGAGC | CCGGTCACGTCTCTTTTTCTC |
| RUNX2 | TGGTTACTGTCATGGCGGGTA | TCTCAGATCGTTGAACCTTGCTA |
| FOXP3 | GTGGCCCGGATGTGAGAAG | GGAGCCCTTGTCGGATGATG |
| BCL6 | GTTGTGGACACTTGCCGGAA | CTCTTCACGAGGAGGCTTGAT |
| LEF1 | AGAACACCCCGATGACGGA | GGCATCATTATGTACCCGGAAT |
| TCF1 | CTGGCTTCTACTCCCTGACCT | ACCAGAACCTAGCATCAAGGA |
| PRDM1 | AACTTCTTGTGTGGTATTGTCGG | CAGTGCTCGGTTGCTTTAGAC |
| RORC | GTGGGGACAAGTCGTCTGG | AGTGCTGGCATCGGTTTCG |

**Supplementary Table 2: Antibody reagents for flow cytometry analysis or sorting**

| Antibody | Clone | Catalog# | Company |
|---|---|---|---|
| **Human antibodies** | | | |
| mouse anti-human CD73 | AD2 | 562430/561254 | BD Biosciences |
| mouse anti-human CD69 | FN50 | 310904 | BioLegend |
| anti-human CD103 | Ber-ACT8 | 350206/350212 | BioLegend |
| anti-human CXCR6 (CD186) | K041E5 | 356010 | BioLegend |
| anti-human CD45 | 2D1 | 368514 | BioLegend |
| anti-human CD45RO | UCHL1 | 304204/304230 | BioLegend |
| anti-human CD45RA | HI100 | 304134 | BioLegend |
| anti-human CD3 | UCHT1 | 300424 | BioLegend |
| anti-human CD4 | RPA-T4 | 300506/300537 | BioLegend |
| anti-human CD4 | OKT4 | 317429 | BioLegend |
| anti-human CD8a | RPA-T8 | 301016/301028 | BioLegend |
| anti-human CD62L | DREG-56 | 304806/304822 | BioLegend |
| anti-human CD127(IL7R$\alpha$) | A019D5 | 351303 | BioLegend |
| anti-human CD117(c-KIT) | 104D2 | 313203 | BioLegend |
| mouse anti-human Ki67 | B56 | 561283 | BD Biosciences |
| anti-human FOXP3 | 206D | 320107 | BioLegend |
| mouse anti-human HLA-A2 | BB7.2 | 561341 | BD Biosciences |
| anti-human IL-2 | MQ1-17H12 | 500322 | BioLegend |
| anti-human TNF$\alpha$ | MAb11 | 502916 | BioLegend |
| anti-human IL-17A | eBio64DEC17 | 12-7179-41 | eBioscience |
| mouse anti-human IFN$\gamma$ | B27 | 562016 | BD Biosciences |
| anti-human IL-21 | 3A3-N2 | 513004 | BioLegend |
| anti-human IL-4 | 8D4-8 | 500703 | BioLegend |
| anti-human/mouse Granzyme B | GB11 | 515406 | BioLegend |
| CD45RO microbeads, human | | 130-046-001 | Miltenyi Biotec |
| **Mouse antibodies** | | | |
| anti-mouse CD103 | 2E7 | 121431 | BioLegend |
| anti-mouse CD186 (CXCR6) | SA051D1 | 151118 | BioLegend |
| anti-mouse CD73 | TY/11.8 | 127212 | BioLegend |
| anti-mouse CD4 | RM4-4 | 116022 | BioLegend |
| anti-mouse TCR$\beta$ chain | H57-597 | 109207 | BioLegend |
| anti-mouse CD69 | H1.2F3 | 104522 | BioLegend |
| anti-mouse CD8a | 53-6.7 | 100706 | BioLegend |
| anti-mouse CD45.1 | A20 | 110714 | BioLegend |
| anti-mouse CD45.2 | 104 | 109808 | BioLegend |
